# PARNET: A CLIP-SEQ-BASED FOUNDATION MODEL FOR RNA SEQUENCE REPRESENTATION LEARNING

**DOI:** 10.64898/2026.08.08.743506

**Authors:** Lambert Moyon, Andreina Tirabassi, Artem Baranowskii, Charlotte Capitanchik, Klara Kuret Hodnik, Leo Wilkinson, Shubankar Londhe, Gabrijela Dumbovic, Julien Gagneur, Jernej Ule, Marc Horlacher, Annalisa Marsico

## Abstract

RNA-binding proteins (RBPs) orchestrate a complex combinatorial regulatory “code” that governs RNA splicing, stability, localization, and translation. Learning the relationship between RNA sequences and these processes is a central challenge in genomics. Foundation models, notably RNA language models, have emerged as the dominant approach, learning general-purpose representations from unlabeled sequence at scale. While RNA language models have demonstrated impressive performance across a broad range of downstream tasks, they generally learn from sequence reconstruction objectives alone, lacking direct connections to the regulatory principles that govern RNA function. Here we introduce Parnet, an RNA foundation model trained directly and exclusively on experimental CLIP-seq data. Parnet is a multi-task foundation model trained end-to-end on 223 eCLIP-seq experiments spanning 150 RBPs to predict base-resolution RBP binding profiles directly from RNA sequence. This CLIP-seq pretraining strategy departs fundamentally from the masked-language-modeling paradigm, anchoring learned RNA representations directly in measured protein–RNA interactions rather than sequence statistics. Parnet substantially outperforms its single-task predecessor RBPNet in binding profile and motif recovery, generalizes to unseen cell types and iCLIP data, and recapitulates position-dependent splicing regulation. Frozen Parnet embeddings, without task-specific fine-tuning, match or exceed the performance of both task-specific tools, as well as larger self-supervised RNA and genomic language models across diverse downstream tasks, including RNA biotype classification, lncRNA chromatin localization, translational efficiency, splice-site recognition, intron retention, and non-coding variant effect prediction. Importantly, Parnet remains mechanistically interpretable, tracing predictions back to the specific RBPs and motifs that drive them. These results establish the RBP interactome as a compact, functionally sufficient, and interpretable basis for foundation model pretraining in RNA biology.

## Background

RNAs are essential regulators of biological processes such as splicing, translation, and post-transcriptional regulation. Their sequence- and structure-dependent interactions with proteins and other nucleic acids make them versatile therapeutic targets and drug modalities. A key layer of post-transcriptional regulation is the combinatorial binding of RNA by RNA-binding proteins (RBPs). The “RBP code” governs how a transcript is processed, stabilized, or degraded within the cell [1]. Although high-throughput experimental methods can provide quantitative measures of RNA metabolic processes [2–6], such as translation, mRNA degradation, splicing and RBP binding transcriptome-wide, they remain labor-intensive and costly. As an alternative, sequence-based supervised machine learning models can learn predictive features directly from DNA and RNA sequences.

Over the past decade, deep learning has transformed genomics, outperforming traditional feature-based methods by learning complex nonlinear patterns directly from raw biological data. Convolutional neural networks (CNNs) capture local sequence dependencies efficiently and remain a core component of models for transcription factor binding, chromatin accessibility, gene expression, and RNA–protein interactions[7–10]. Their limited receptive field motivated recurrent architectures such as RNNs and LSTMs, which model longer-range dependencies [11]. Recent advances have enabled accurate prediction of splicing (SpliceAI [12], DeepSplice [13], MMSplice [14]), translation efficiency (Optimus 5-Prime [15], FramePool [16], RiboNN [17]), and transcript stability (Saluki [18]) directly from sequence. For RBP–RNA interactions specifically, models have progressed from sequence-based classifiers trained on either *in vitro* data[7] or high-confidence in-vivo CLIP-seq peaks toward richer frameworks incorporating sequence conservation, genomic context, RNA secondary structure, and proximity to splice or transcription start sites ([19], [20], [10],[21]), including models such as PRISMNet [22] and iDeepS [23]. More recently, Transformer-based architectures have modeled post-transcriptional gene regulation more broadly, capturing long-range dependencies through attention rather than recurrence, and have been applied to various tasks spanning RBP–RNA binding [24, 25], miRNA targeting [26], and splicing regulation [27, 28], Beyond direct prediction, sequence-based deep learning models also enable *in silico* perturbations that estimate variant effects on regulatory processes and link genetic variants to functional outcomes, which is critical for variant prioritization and QTL mapping, especially as most GWAS variants lie in non-coding regions [8, 21, 29–32].

Orthogonal to these advances, the prediction target itself has shifted. Deep learning models such as BPNet [33], chromBPNet [34], Enformer [35], and Borzoi [36] learn base-resolution regulatory signals directly from DNA, moving beyond binary classification toward end-to-end sequence-to-signal frameworks. Similar architectures have been adapted for post-transcriptional regulation, to predict continuous CLIP-seq read coverage [37]. On this line, we have previously introduced RBPNet [32], a sequence-to-function, high-resolution sequence-based model trained on individual CLIP-seq experiments to predict base-resolution RBP–RNA binding profiles while correcting for background signal noise. Trained at sufficient scale, such profile prediction yields representations that transfer beyond the task they were fitted on. This is the defining property of foundation models: they are trained on sufficiently large and diverse corpora to learn general-purpose internal representations that transfer effectively across a wide range of downstream tasks, often matching or surpassing task-specific models while requiring only minimal fine-tuning. In genomics and RNA biology, this has largely been pursued through self-supervision: inspired by BERT and GPT, genomic foundation models such as Nucleotide Transformer [38], Evo2 [39], and Orthrus [40] use self-supervised objectives on large unlabeled sequence corpora to learn context-aware representations that enable accurate prediction with minimal fine-tuning. RNA-specific foundation models [41–44] extend this paradigm to holistic modeling of RNA function. However, reconstruction-based self-supervised objectives optimize a proxy for sequence likelihood rather than for regulatory function, and thus may not efficiently capture the sequence-based functional features most relevant to RNA processes. This motivates complementary biologically-grounded pretraining strategies. Orthrus, for instance, uses contrastive pretraining on matured mRNAs, maximizing similarity between splicing isoforms and evolutionarily related transcripts across 400+ mammalian species to focus representations on conserved, information-rich regions ([40]). Supervised models trained directly on experimental measurements, such as Borzoi ([36]) and BigRNA ([45]) retain foundation-like versatility while remaining anchored to functional data.

Here, we postulate that knowing the binding partners of an RNA provides direct insight into both how the RNA is processed and its potential role in downstream processes, and that a model densely representing an RNA’s interactome can serve as a foundation for a wide range of additional tasks. Such a model would be extending the philosophy of Borzoi and BigRNA into the post-transcriptional domain, by grounding RNA representation in the RBP interactome. Building on our previous single-task model RBPNet for predicting RBP-RNA footprints, we introduce Parnet, an RNA foundation model trained end-to-end to predict RBP binding profiles directly from raw RNA sequences derived from CLIP-seq data.

Parnet first extends RBPNet to a multi-task framework, simultaneously learning multiple RBP binding profiles, and doing so benefits from the genomic signals being correlated across related processes, including co-binding RBP patterns. Furthermore, Parnet functions as a foundation model for learning downstream RNA processes and properties.

We show that Parnet markedly improves the state-of-the-art, both in terms of the accuracy of single-nucleotide binding profile prediction, as well as the recovery of genuine RBP sequence motifs. Beyond predictive accuracy, Parnet captures the regulatory logic of RNA processing, faithfully recapitulating functional splicing events—including exon enhancement and silencing, and their associated regulatory mechanisms, across multiple RBP datasets. Owing to its supervised training on RBP-binding data, Parnet learns a rich, functional representation of RNA sequences that encodes the determinants governing how an RNA is regulated. Despite having a much smaller size than most current RNA foundation models (21 million parameters), Parnet reaches performances equivalent to, or higher than, existing RNA and genomic foundation models, as well as task-specific deep learning tools. This is true across a range of downstream tasks including RNA functional classification, splicing, intron retention, mRNA translation prediction, even with minimal or no fine-tuning, and improves prioritization of splicing and pathogenic variants. The supervised training of Parnet, together with explainable AI analyses, can identify which RBPs drive predictions of specific RNA properties or processes, and can dissect the regulatory consequences of non-coding variants in terms of RBP binding perturbation.

Together, our Parnet model demonstrates that experimentally grounded, RBP interactome-aware pretraining is a powerful and interpretable foundation for RNA biology, bridging the gap between single-task binding prediction and integrated modeling of post-transcriptional regulation.

## Results

### Parnet: From CLIP-seq–Based RBP Binding to RNA Representation Learning

We developed Parnet, a multi-task sequence-to-function model designed to predict base-resolution RNA-binding protein (RBP) binding profiles trained end-to-end on raw eCLIP-seq data. Conceptually, Parnet shifts away from traditional single-task models by simultaneously learning on 223 different eCLIP experiments from ENCODE. This multi-task approach allows the model to capture the complex, combinatorial RBP-binding patterns that govern post-transcriptional regulation.

As illustrated in Figure 1a, the model architecture, building upon our previous single-task RBPNet model, begins with the processing of human gene annotations into fixed-length tiles, which are then one-hot encoded and passed through a series of 1D convolutional layers and residual blocks. This backbone converts the raw sequence into highdimensional nucleotide-wise embeddings. From this shared representation, the model branches into 223 task-specific output heads, each corresponding to a unique eCLIP experiment. To ensure specificity, similarly to RBPNet, each head utilizes a “total track” that is modeled as a mixture of two components: a protein-specific “target track” and an experimental “control track” (Size-Matched Input). An additional penalty in the Loss function (see Methods) incentivizes the model to disentangle genuine biological binding signals from experimental biases. Figure 1b shows Parnet predictions in inference mode for a representative transcript and for 3 RBPs selected based on their high bindingsite density (ENCODE NarrowPeaks), illustrating the model’s ability to faithfully recover complex protein–RNA interaction landscapes. The predicted profiles closely match multiple layers of experimental evidence, including raw read-start counts, crosslink sites identified by the single-nucleotide resolution crosslink detection tool PureCLIP, and replicate-specific ENCODE NarrowPeaks.

**Figure 1:**
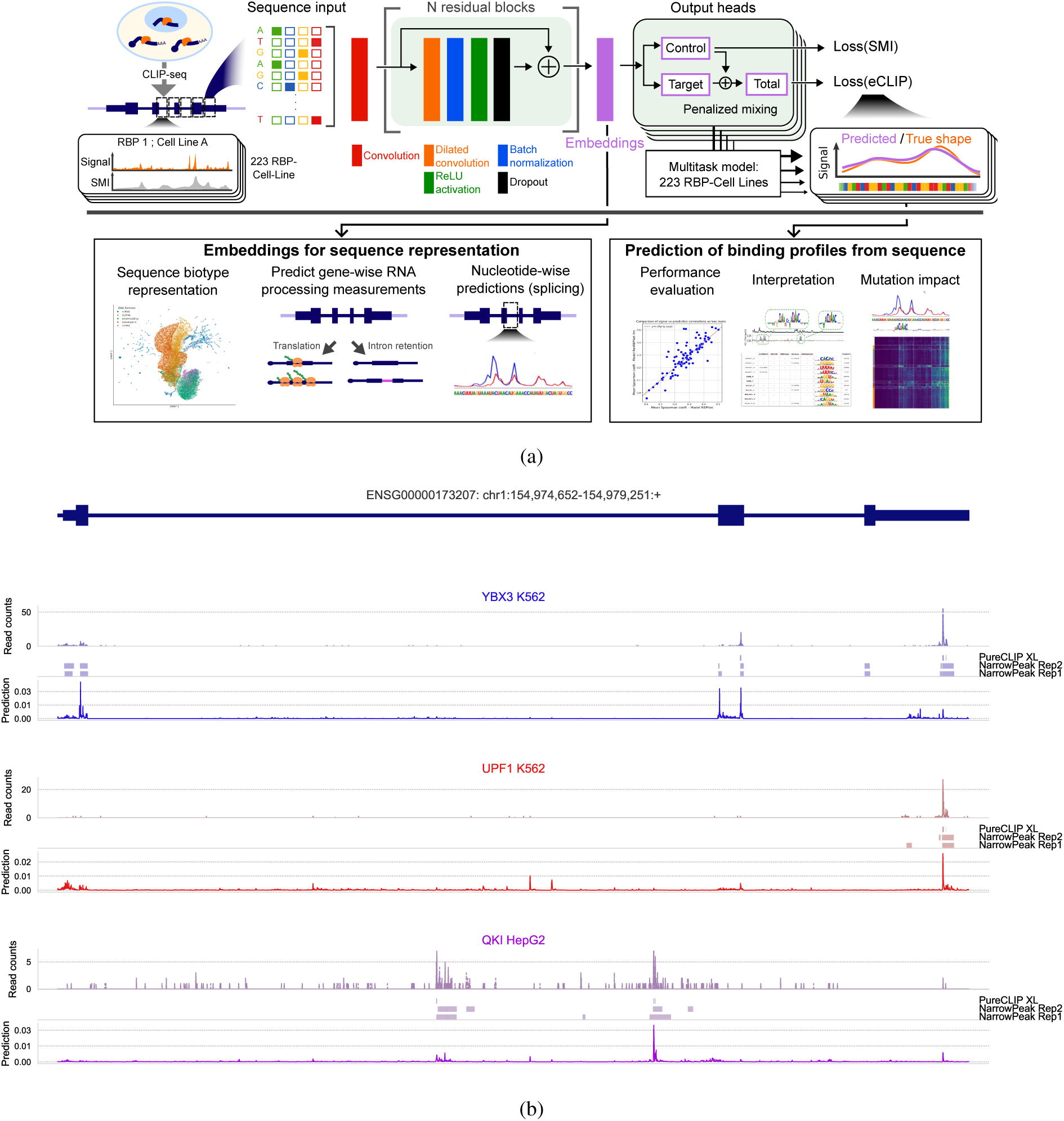
Overview of the Parnet model. **A** Schematic overview of data processing, PARNET training, and downstream applications. Gene annotations are segmented into fixed-length tiles, whose one-hot encoded RNA sequences are passed through a 1D convolution followed by stacked residual blocks. The final block outputs are used for multi-task prediction of *target* and *control* probability tracks via transposed convolution, with the *total* binding signal modeled as a penalized additive mixture of both. Each dataset is assigned a dedicated prediction head, enabling joint prediction across RBPs. Predictions support performance evaluation, comparison to single-task models, binding interpretation via sequence motif retrieval, and *in silico* non-coding mutation scoring. Additionally, outputs of the final residual block are used as nucleotide embeddings, enabling RNA biotype classification, prediction of transcript-level regulatory features (e.g., splicing, intron retention, translation), as well as nucleotide-wise classification os splice sites. **B** Example of transcript-wide predictions by PARNET. Three RBPs with the largest number of NarrowPeaks over the transcripts are displayed, showing from top to bottom the measured read-start counts from raw eCLIP experiments, cross-link sites as extracted by PureCLIP, NarrowPeaks from replicate 1 and 2 as computed in ENCODE, and the Parnet predicted profiles.

Beyond predicting RBP-binding profiles, the model has learned nucleotide-wise embeddings that are inherently tied to protein-RNA interaction logic, and can be extracted and repurposed for a wide range of downstream tasks. In the following, we demonstrate the capability of those embeddings to classify genomic regions and RNA biotype, including different functional classes of long-non coding RNAs, predict mRNA translational efficiency, splice sites and intron retention events, and score the effects of non-coding variants, highlighting Parnet’s versatility as a foundational tool for RNA biology.

### Parnet improves base-resolution RBP binding’s prediction accuracy over single-task RBPNet and generalizes to iCLIP dataset

We first assessed Parnet’s ability to predict RBP binding profiles on unseen RNA transcripts (i.e. regions excluded from training; see Methods), compared to single-task RBPNet. For a fair comparison, we retrained RBPNet on longer input sequences, mirroring Parnet’s training sequence length (600 nucleotides) and extended it to all CLIP-seq datasets used by Parnet, beyond the original 103 ([46]; see Methods). Parnet showed strong performance on held-out transcripts, achieving Pearson correlations (r) ranging from 0.17 to 0.70 across RBPs (*mean* = 0.48); Figure 2a), and outperforming all single RBPNet models. Overall, Parnet improves prediction accuracy, with an average increase of 35%in Pearson correlation (Figure 2a) and 19% in Spearman correlation (Figure 1a) between predicted and observed read-start count profiles relative to single-task RBPNet models.

**Figure 2:**
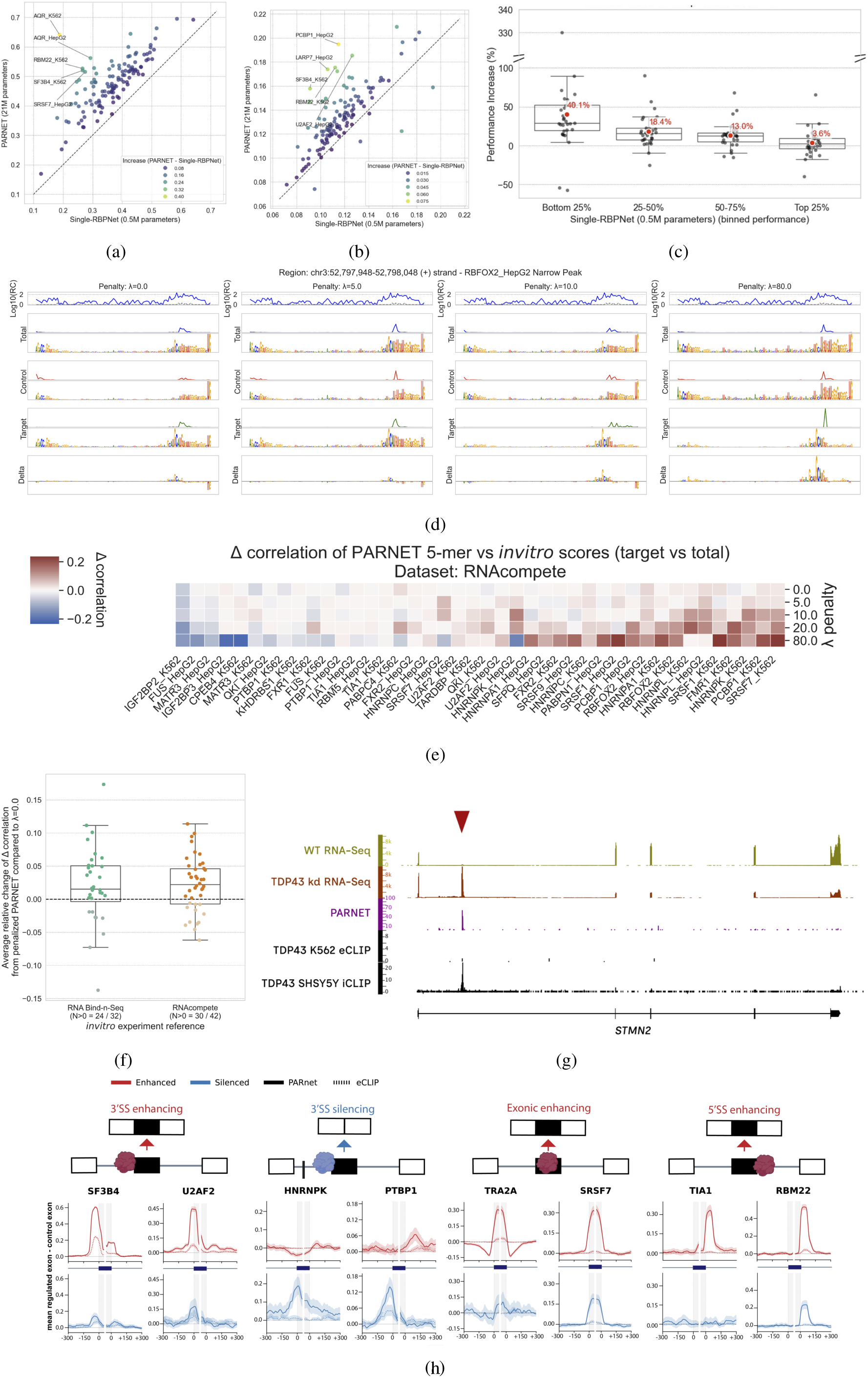
Performance evaluation of the PARNET model. **a.** Comparison of Parnet and reference single-RBPNet performance, measured by the Pearson correlation between predicted profiles and observed read-start counts across transcriptomic regions on the test-set chromosomes. **b.** Comparison of Parnet and reference single-RBPNet performance, measured by the auPRC of scored ENCODE NarrowPeaks on the test-set chromosomes. **c.** Relative performance improvement of Parnet over the reference single-RBPNet model, stratified by quantiles of reference model performance (Spearman correlation between predicted profiles and observed read-start counts). **d.** Effect of the penalty parameter (*λ*) on Parnet signal denoising and motif recovery for a representative RBFOX2-bound region. Panels correspond to *λ* = 0 (no penalty), 5, 10, and 80. Subpanels (top to bottom) show log_10_ read counts and original CLIP signal; total Parnet prediction and its integrated gradients motif; predicted control track and motif; predicted target track and motif; and the integrated gradients map highlighting sequence features distinguishing target from control. Increasing *λ* progressively denoises the target signal and recovers a motif that more closely resembles the canonical RBFOX2 binding motif. **e.** Difference (Δ) in correlation between *in vitro* 5-mer scores (from RNACompete assays) and Parnet feature importance across increasing penalty. For each RBP, Δ is computed as the difference between correlations obtained from the denoised target and total prediction tracks. Higher Δ indicates improved agreement between *in vivo* and *in vitro* motifs and more faithful recovery of genuine target motifs. **f.** Mean Δ across increasing penalty for RNA Bind-n-Seq and RNAcompete experiments. Positive values indicate stronger agreement between *in vitro* scores and Parnet feature importance from the denoised target track. **g.** Comparison of experimentally measured TDP-43 binding profiles across the full STMN2 transcript. Parnet recapitulates the iCLIP binding profile, whereas the corresponding signal is absent from eCLIP. **h.**Exon–intron meta-profiles of RBP binding derived from eCLIP signal (dotted lines) and Parnet predictions (solid lines) for four classes of splicing changes following knockdown of eight selected RBPs.

We next evaluated the agreement between Parnet predictions and experimentally defined RBP binding sites using both peak-level and single-nucleotide annotations. Specifically, we compared Parnet predicted probabilities with ENCODE NarrowPeaks (called by CLIPper [47] and with crosslink (CL) sites identified by PureCLIP [48] (see Methods). Across RBPs, Parnet consistently outperforms single-task RBPNet models in both peak-level and single-nucleotide evaluations. For ENCODE NarrowPeaks, Parnet achieves higher AUPRC and AUROC for nearly all RBPs, with average gains of 21% and 1%, respectively, and maximum AUPRC improvements reaching up to 148%. Similar trends are observed for PureCLIP-defined crosslink sites, with average increases of 40% (auPRC) and 1% (AUROC), again with substantial gains for individual RBPs.

Model performance is influenced by experimental data quality, for which we considered two proxies: the number of called peaks and the fraction of uniquely aligned (non-PCR duplicate) reads. Consistent with our previous benchmark study ([49]), we observe a positive correlation between predictive performance and the number of detected peaks, supporting the notion that higher signal-to-noise facilitates learning. For both proxies, the association with performance is strongest at the low end and flattens for well-covered datasets. Despite this, Parnet yields the largest improvements for RBPs with lower baseline performance in single-task models. Specifically, RBPs in the bottom 25% quantile of RBPNet performance exhibit an average gain of 40% in Spearman correlation, significantly higher when compared to only 3.6% for RBPs in the top quantile. These results highlight Parnet’s ability to robustly improve prediction accuracy, particularly for challenging, lower-quality datasets. While these gains is likely a consequence of Parnet’s multi-task design, where RBPs with similar binding preferences, particularly in lower-quality datasets, benefit from shared representations, it is important to note that Parnet has substantially higher capacity (21 Million parameters) than RBPNet (0.5 Million) and a wider sequence context window to learn from (see Methods). Thus, part of the performance improvement may also stem from the increased model complexity.

### Parnet’s penalty λ enhances recovery of specific RBP sequence signals and *in vitro* motifs

A key application in RNA–RBP interaction modeling is to identify sequence features that govern recognition of the RNA by a given RBP. Attribution methods such as Integrated Gradients (IGs) highlight sequence elements that drive predictions, capturing RBP-specific binding preferences that can be summarized as sequence motifs (see Methods). Here, wanted to assess the impact of the *λ* penalty, which regulates the mixing of control and target signals, on motif CLIP signal motif specificity. Increasing *λ* forces greater reliance on the control track. This denoises the predicted eCLIP signal and is expected to enhance motif specificity.

To illustrate the effect of (*λ*) on the retrieved binding motif, Figure 7c shows IGs attribution maps for RBFOX2 computed with respect to the total, control, and target tracks at different values of *λ* (*λ* = 0, 2.5, 10, and 80), together with the corresponding predicted signals in a NarrowPeak 100 nucleotides region identified in the gene ITIH3 (blue: total, red: control, green: target). At low *λ* values, the IG maps recover a composite motif consisting of the canonical RBFOX2 motif together with a G-rich sequence pattern, previously described by us and others [32, 50, 51] as background signal in both control and target signal, as well as predicted total signal. Increasing *λ*, particularly from *λ* = 10 onward, results in substantially higher specificity of the target track toward the consensus RBFOX2 motif, while the G-rich pattern becomes predominantly associated with the control track. This suggests that stronger penalization of the target–control mixing coefficient enables more effective separation of genuine binding specificity from experimental background noise, qualitatively illustrating the effect of *λ* on the specificity of motifs recovered from the predicted signal track. A similar trend is observed for QKI, where higher *λ* values lead to progressive denoising of the canonical QKI motif in the target track, while the putative background U-rich motif (UUUU) becomes largely confined to the control track (Supplementary Figure 2b).

In our previous model, RBPNet, we introduced an orthogonal validation strategy to assess whether learned sequence features reflect genuine RBP binding preferences, by comparing inferred motifs to *in vitro* protein–RNA interaction data, namely RNA Bind-n-Seq [52] and RNAcompete [53]. In particular, we demonstrated that modeling the control signal as an auxiliary task improves the specificity of motifs derived from CLIP data. Here, we extend this analysis with Parnet by incorporating updated RNAcompete datasets, which cover a broader range of RBPs [54]. We find that Parnet-derived RBP sequence preferences (5-mers) extracted from the denoised target track show increasing agreement with in vitro motifs (RNAcompete or RNA Bind-n-Seq) as the *λ* penalty is raised from 0 to 5, 10, 20, and up to 80, compared to motifs derived from the unpenalized total track (quantified as Δ correlation in Figure 2e and Supplementary Figure 2a; higher values indicate stronger in vitro concordance). This indicates that denoising the target track yields motifs that more faithfully recapitulate intrinsic sequence preferences. Consistently, pooling across all Parnet models and RBPs with matched *in vitro* data, this trend holds on average for both assay types (Figure 2f. However, overly strong penalization (e.g. *λ* = 80) reduces agreement, likely due to loss of motif signal in the total track caused by excessive shrinkage toward the control. Overall, *λ* = 10 provides the best trade-off, yielding a consistent improvement in correlation across RBPs while maximizing coverage, and is therefore used for downstream analyses.

### Parnet recovers position-dependent splicing regulation and binding in unseen cell lines

RNA-binding proteins (RBPs) can either enhance or repress the inclusion of cassette exons depending on where they bind relative to the splice sites. This positional regulation influences which transcript isoforms are produced, and these splicing outcomes can be reconstructed and quantified from RNA-seq data. Because ParNet does not use RNA-seq data as input, it cannot directly learn these regulatory relationships from observed splicing outcomes. Instead, in the absence of RNA-seq measurements that quantify exon regulation, we sought to determine whether ParNet can nevertheless identify functionally meaningful RBP binding sites—that is, binding sites whose presence is associated with changes in exon inclusion or skipping. We did this by performing an RNA splicing map analysis. RNA splicing maps are a way to integrate transcriptome-wide RBP binding data with splicing profiles to reveal such position-dependent regulatory effects ([55]). They summarize RBP binding patterns around alternatively spliced exons by aggregating binding signals across exons that are enhanced, silenced, or unaffected by a given RBP. Resulting RNA splicing maps often identify distinct positional binding principles that underlie positive or negative activity of specific RBPs on exon inclusion.

While such maps are usually constructed from CLIP experimental signal, here we aimed at assessing whether Parnet predictions can correctly capture the position-specific binding patterns. Therefore, we built RNA maps by integrating RBP knockdown RNA-Seq data either with Parnet predictions or CLIP experimental signal for comparison. If, for example, a particular RBP is known to enhance or silence a regulated exon depending on where it binds, we evaluated whether Parnet assigns strong predicted binding signals to the corresponding regulatory regions around those exons.

Parnet faithfully recovered signal at the 3’SS of exons enhanced by core spliceosomal proteins SF3B4 and U2AF2; the exonic enhancing binding of serine-arginine rich (SR) proteins TRA2A and SRSF7; and the 5’SS-enhancing binding of TIA1 and RBM22 (Figure 2h). Further, the model predicted 3’SS binding at exons silenced by PTBP1 and HNRNPK. Taken together, we demonstrate that Parnet efficiently predicts functionally important binding sites that distinguish between the different modes of regulation of diverse RBPs (Supplementary Figure 3).

A further motivation in developing Parnet is to enable prediction of RBP binding in unseen cell types, which is crucial for advancing our understanding of human disease. We highlight a representative example of this in the *Stathmin2* gene (*STMN2*), a key therapeutic target in the treatment of the neuro-degenerative disease Amyotrophic Lateral Sclerosis (ALS). In patients with ALS, TDP-43 becomes mis-localized to cytoplasmic aggregates, leading to de-repression of cryptic exons in neurons. Despite being trained on TDP-43 K562 eCLIP data where *STMN2* is not expressed, Parnet successfully recovers TDP-43 binding at the cryptic exon, which is further validated by TDP-43 iCLIP signal in SH-SY5Y cells (Figure 2g).

### Parnet embeddings capture functional RNA properties

Once pre-trained, Parnet’s output embeddings provide a powerful sequence representation that encodes functional RNA properties relevant to RNA–protein interactions. We first leverage these embeddings to visualize and cluster RNA sequences in a biologically meaningful space. Afterwards, we use them as inputs for downstream prediction tasks and demonstrate that they improve performance by incorporating information about RBP-binding potential directly into the representation. We assessed the extent to which Parnet embeddings capture sequence features associated with four major genomic elements (CDS, 3*^′^* UTRs, 5*^′^* UTRs, and introns), as well as five RNA families, including protein-coding messenger RNAs (mRNAs), long non-coding RNAs (lncRNAs), microRNAs (miRNAs), pseudogenes, and the broader class of ncRNAs encompassing multiple non-coding RNA subfamilies (Figure 3a).

**Figure 3:**
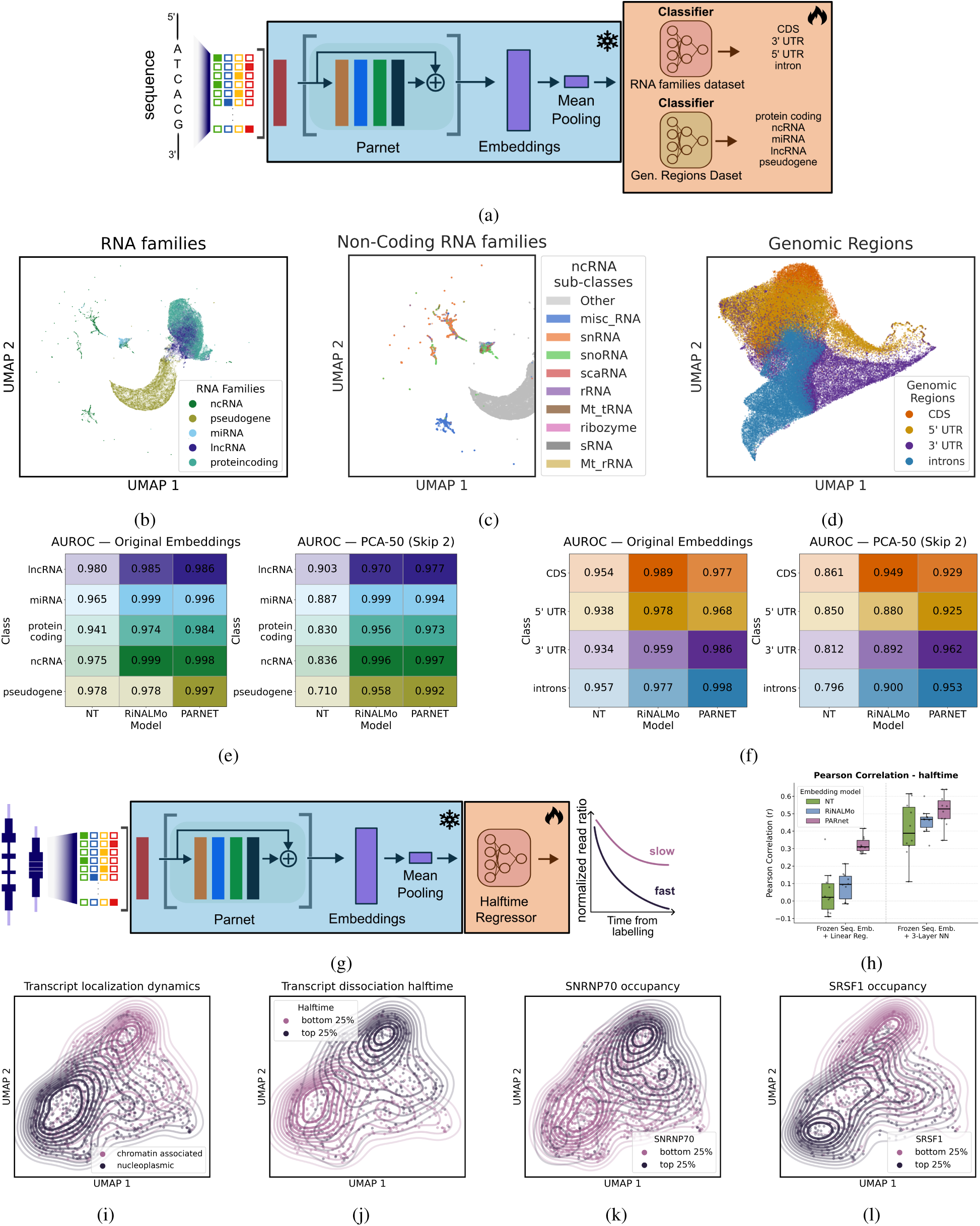
Parnet for gene representations. Figure 3. (a) Overview of downstream genomic prediction tasks performed using frozen Parnet embeddings, including genomic region and RNA family classification. (b) Two-dimensional UMAP projection of Parnet embeddings colored by RNA family. (c) Zoomed-in view of the Parnet embedding space highlighting small RNA subtypes. (d) UMAP projection of Parnet embeddings colored by genomic region annotation. (e) Left: AUROC performance of lightweight classifiers trained on the original embeddings from Parnet, Nucleotide Transformer, and RINALMO for the RNA family classification task. Right: performance on the same task after removing the first two principal components from the embeddings. (f) Left: AUROC performance of lightweight classifiers trained on the original embeddings from Parnet, Nucleotide Transformer, and RINALMO for the genomic region classification task. Right: performance on the same task after removing the first two principal components from the embeddings. (g) Overview of the downstream lncRNA half-life prediction task using frozen Parnet embeddings. (h) Predictive performance (Spearman correlation) of Parnet compared to RINALMO and Nucleotide Transformer using a linear regression model (left) and a three-layer neural network (right) trained on frozen embeddings. (i–j) Two-dimensional UMAP projections of the Parnet embedding space colored by lncRNA localization dynamics (i) and dissociation half-life (j) respectively. Contour lines indicate the upper and lower 25% quantiles of the corresponding distributions. (k) Parnet embedding space colored by SNRNP70 occupancy (ENCODE peaks), with contour lines marking the upper and lower 25% quantiles of the occupancy distribution. (l) Parnet embedding space colored by SRSF1 occupancy (ENCODE peaks), with contour lines indicating the upper and lower 25% quantiles of the occupancy distribution.

Unsupervised visualization of the Parnet embeddings reveals that the model learns to distinguish sequences originating from coding, intronic, and untranslated regions (UTRs), as well as different RNA families and ncRNA subfamilies, albeit to varying degrees (Figure 3b–d), similarly to Nucleotide Transformer [38] and RiNALMo [42] (Supplementary Figure 3b, Supplementary Figure 3c). To further quantify this information, we trained lightweight classifiers on top of globally pooled embeddings (Figure 3a; see Methods), achieving high classification accuracies across all categories (Figure 3f, Figure 3g). Notably, Parnet performs on par with (or surpasses) existing DNA and RNA language models, including RiNALMo and Nucleotide Transformer, across both tasks.

Given the strong performance of both Parnet and previously proposed RNA/DNA language models on this classification task, we next investigated which features these models capture within their embeddings, and whether part of their predictive power could arise from underlying dataset biases. To address this, we first examined whether Parnet embeddings for the RNA family classification task exhibited gradients associated with potentially confounding features, including sequence length, GC content, and RNA expression levels derived from ENCODE RNA-seq data in both K562 and HepG2 cells. This analysis aimed to determine whether the model learns informative biological representations beyond such trivial features. Visual inspection of the embedding space revealed a noticeable dependence on sequence length, as expected, with pseudogenes and small RNAs separating from the remaining classes due to their substantially longer or shorter transcript lengths, respectively (Figure 3g). In contrast, we observed no clear bias associated with RNA expression levels, an important potential confounder given that the model is trained on CLIP-seq data, nor with GC content (Supplementary Figure 3f, Supplementary Figure 3h, Supplementary Figure 3i). We extended this analysis to additional foundation models, including Nucleotide Transformer and RiNALMo, and quantitatively assessed the relationship between embedding dimensions and potential biases. To this end, we applied PCA to the learned embeddings and examined the correlation between the principal components and features such as sequence length and GC content. This analysis revealed that the leading principal components of both Nucleotide Transformer, and, to a lesser extent, Parnet, were significantly correlated with sequence length, particularly in the genomic region and RNA family classification tasks. In contrast, RINALMO showed a stronger association with GC content. To further evaluate the contribution of these biases to downstream performance, we removed the first two principal components and re-projected the sequences into the residual embedding space. While this had little effect on Parnet’s classification accuracy, it substantially reduced the performance of the other language models, in some cases by more than 10% (Supplementary Figure 3o, Supplementary Figure **??**). This increased the performance gap in favor of Parnet, suggesting that existing RNA/DNA language models may be more affected by global sequence biases, whereas Parnet captures more functionally relevant sequence features, likely due to its training on RNA–protein interaction landscapes.

Having demonstrated that Parnet embeddings can capture the characteristics of distinct genomic elements and RNA families in an unsupervised manner, we next explored their utility in more complex functional genomics tasks. As a first application, we focused on classifying lncRNAs as either nucleoplasmic or chromatin-associated. In our previous work [56], we developed chrTT-seq (chromatin transient transcriptome sequencing), a framework combining pulse-chase metabolic labeling with cellular fractionation to profile the subnuclear kinetics and chromatin dissociation dynamics of newly transcribed RNAs. By mathematically modeling RNA release rates, transcripts could be classified according to their chromatin association half-lives, distinguishing rapidly released nucleoplasmic RNAs from chromatin-tethered species. A key finding from that study was that enhancer-derived lncRNAs enriched for enhancer-associated chromatin marks are released into the nucleoplasm particularly rapidly. Using classical machine learning approaches, we further showed that molecular features such as splicing efficiency and binding probabilities for specific RNA-binding proteins (RBPs) are major determinants of whether an RNA remains chromatin-associated or is efficiently released.

We assessed whether Parnet embeddings computed from full-length transcripts could predict lncRNA chromatin association half-lives in a zero-shot setting using data from the chrTT-seq study. To this end, we trained either a simple linear regression model or a three-layer neural network on top of frozen Parnet embeddings (Figure 3h). Parnet achieved strong predictive performance, reaching a Pearson correlation of 0.33 with linear regression and 0.54 with the neural network between predicted and experimentally measured half-lives. Importantly, Parnet substantially outperformed both Nucleotide Transformer and RiNALMo on this task (3i). The learned embedding space naturally separated rapidly released, nucleoplasmic lncRNAs from chromatin-tethered transcripts with long chromatin half-lives (Figure (**??**, 3k)), demonstrating that Parnet captures biologically meaningful features associated with RNA localization and dissociation dynamics. Furthermore, interrogation of the embedding space by overlaying RBP occupancy and chromatin-associated features recapitulated several of the main determinants identified in the original [56]. In particular, chromatin-tethered lncRNAs clustered with enrichment for the spliceosomal factor SNRNP70 (3l), whereas rapidly dissociating lncRNAs were associated with SRSF splicing factors, enhancer-associated chromatin marks, and the transcription termination factor XRN2 (Supp. Fig. 5g) Together, these results highlight the ability of Parnet embeddings to capture higher-order functional properties of RNAs and to generalize to complex downstream tasks beyond direct RBP-binding prediction.

### Fine-Tuning the Parnet model to predict and interpret RNA processes Prediction of mRNA Translational Efficiency (TE)

mRNA translation is the process by which ribosomes decode an mRNA sequence to synthesize proteins. Translational Efficiency (TE) quantifies how effectively a transcript is translated—typically measured in different cell lines as the ratio of ribosome occupancy from Ribo-seq experiments [57] to mRNA abundance (RNA-seq). We evaluated whether Parnet embeddings capture sequence features relevant for predicting TE by training downstream models on frozen full-length transcript embeddings (Fig. 4). We benchmarked two approaches: a linear regression model and a two-layer neural network trained on Parnet embeddings (Supplementary Figure 4a), across four independent datasets spanning HEK293 cells, muscle tissue, and PC3 cells. Since known determinants of TE, such as GC content and transcript length, are also potentially encoded in RNA language model representations (Fig. 4a), and are also predictive factors of mRNA TE ([58]), we compared Parnet-based models against a baseline linear regression model using only GC content and transcript length as input features. This allowed us to assess whether Parnet embeddings capture additional regulatory information beyond these basic transcript-level properties.

**Figure 4:**
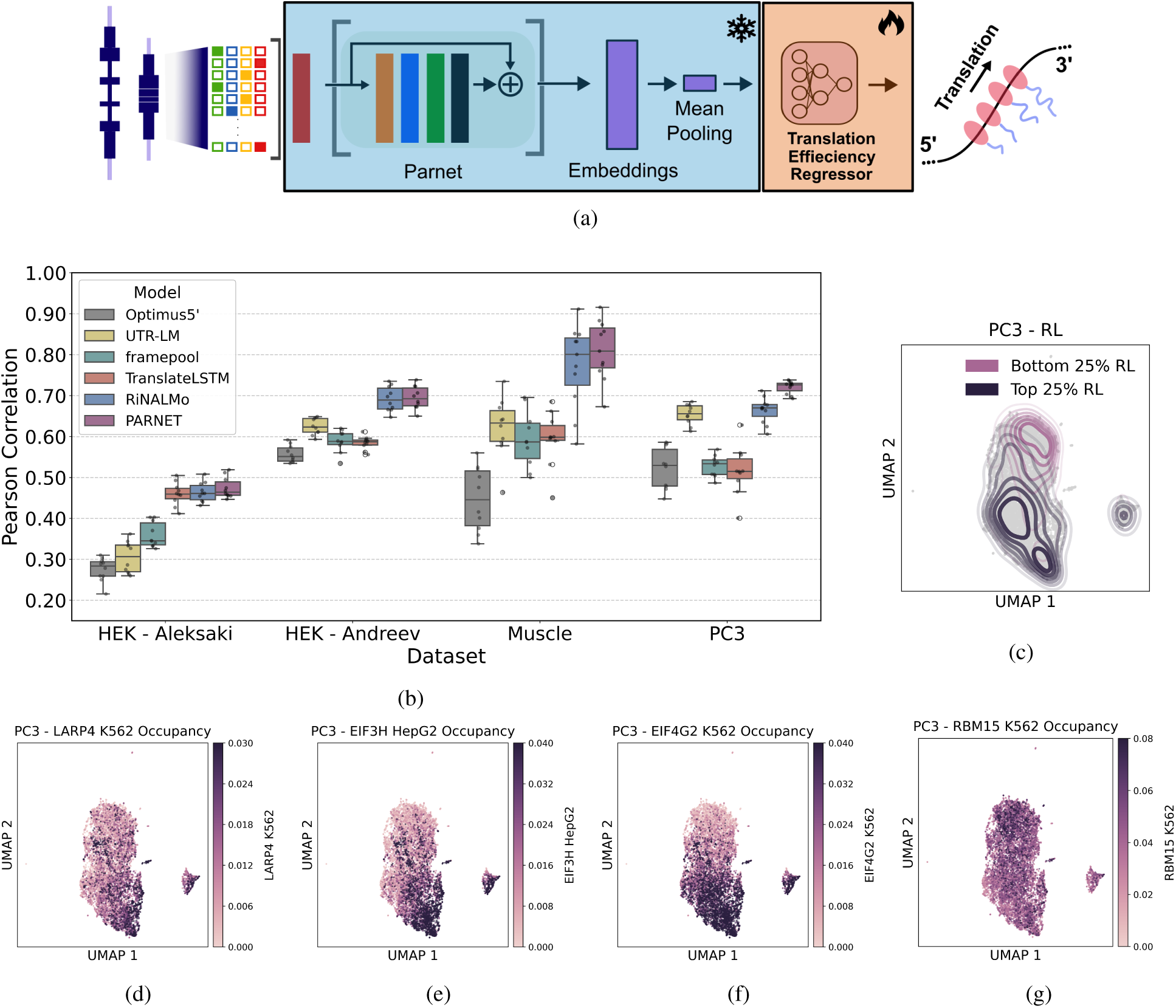
(a) Overview of the downstream translational efficiency (TE) prediction task using ribosome loading data and frozen Parnet embeddings. (b) Two-dimensional UMAP projection of the Parnet embedding space colored by translational efficiency in the PC3 dataset. Contour lines indicate the upper and lower 25% quantiles of the TE distribution. (c) Predictive performance (Pearson correlation) of Parnet compared with RINALMO, TranslateLSTM, UTR-LM, FramePool, and Optimus5’ (100 nt). Boxplots show the distribution of performance across the 10 cross-validation folds for a two-layer neural network regressor trained on frozen embeddings to predict TE. (d-g) Two-dimensional UMAP projections of the Parnet embedding space colored by occupancy of the RBPs LARP4 (d), EIF3H (e), EIF4G2 (f) and RMB15 (g).

Projection of Parnet embeddings into a two-dimensional UMAP space revealed a clear organization of transcripts according to TE values, with stronger separation between transcripts in the upper and lower quartiles of the TE distribution (Fig. 4c). A similar pattern was observed in the HEK293 dataset from [59] using mean-pooled Parnet embeddings (**??**), further supporting that Parnet representations encode information associated with translational regulation.

We also benchmarked Parnet against five state-of-the-art methods: Optimus 5*^′^*, one of the earliest deep learning models to predict translational efficiency from 5*^′^* UTR sequence features using CNNs trained on high-throughput reporter assays [60]; FramePool, an extension of Optimus 5*^′^* that handles variable-length UTRs by aggregating features across all possible reading frames via convolution and pooling architectures [16]; TranslateLSTM, an LSTM-based model that predicts translational efficiency from mRNA sequence context, incorporating UTR and coding-region features as well as handcrafted descriptors such as minimum free energy, k-mer composition, GC content, transcript length and others [61]; and UTR-LM, a language-model approach that learns representations of 5*^′^* UTR sequences to infer translational efficiency from sequence context [62]. These four methods are specifically designed for mRNA translation prediction. In addition, we included the general-purpose RNA language model RiNALmo that has already demonstrated strong performance on this task [42]. In cross-validation experiments, Parnet consistently outperformed competing methods across all four datasets and three cell types based on Pearson correlation (Fig. 4d). The Parnet-based model using a two-layer neural network achieved the highest predictive performance, outperforming both the GC content and transcript length baseline model and the linear regression model trained on Parnet embeddings (Supp. Fig. 6a). While performing comparable to RiNALMo in HEK293 cells, Parnet achieved higher accuracy in muscle cells (0.82 vs. 0.78) and more in PC3 cells (0.73 vs. 0.67).

Because Parnet is trained on CLIP-seq data, its embedding space can be interrogated by projecting training sequences into a two-dimensional UMAP representation and coloring transcripts according to “RBP occupancy”, defined, for each transcript, as the number of peaks called from Parnet’s predictions per transcript (see Method). This provides an indirect but intuitive way to identify which RBPs may underlie a given downstream prediction. Although qualitative in nature, this approach offers interpretability in settings where attribution methods such as Integrated Gradients cannot be readily applied.

Using this strategy, we identified several RBPs strongly associated with the separation between high- and low-TE transcripts in the PC3 dataset (Fig. 4d). Among the most prominent were LARP4, which binds poly(A) tails and promotes efficient ribosome loading and translation (Fig. 4e); EIF3H, a core component of the eIF3 complex involved in stabilizing ribosome recruitment at the 5*^′^* UTR (Fig. 4f); EIF4G2, a non-canonical translation initiation scaffold linking mRNAs to the ribosome (Fig. 4g), and RBM15 (Fig.**??**), a key regulator of m6A RNA modification and processing, which may indirectly influence TE through effects on RNA stability. Its enrichment in the low-TE region of the Parnet embedding space is consistent with the established role of m6A regulation in modulating RNA half-life. From inspection of the embedding space, a positive association of EIF4G2 and LARP4 with high-TE transcripts is also observed in the HEK293 Aleksaki dataset (Supp. Fig. 6c), together with LIN28 (Supp. Fig. 6e), which is interestingly an indirect positive regulator of translation through inhibition of let-7 miRNA biogenesis. In HEK293, we also recover a similar pattern of indirect regulation with XRN2 (Supp. Fig. 6f), an RBP involved in mRNA stability.

### Prediction of donor and acceptor splice sites for human and other species

We next leveraged Parnet for the classification of splice sites versus non-splice sites using a widely used benchmark derived from the G3PO+ dataset[63]. This dataset contains experimentally curated splice-site sequences from 148 eukaryotic organisms, including human, with positive and negative sets constructed by sampling from exon and intron regions of the corresponding genomic sequences. The test set comprises four species that are not seen during training. The same benchmark was also used to evaluate the general-purpose RNA language model RiNALMo and the deep learning model Spliceator. Instead of binary classification (donor vs. non-splice and acceptor vs. non-splice, as done in [63] and [42]), we considered a more challenging ternary classification setting, in which the model must simultaneously distinguish donor sites, acceptor sites, and non-splice sequences (Fig. 5a). This task is more difficult as it requires joint discrimination between both splice-site classes in addition to negative sequences. For Parnet, we used frozen full-length embeddings extracted from the pretrained encoder and trained a shallow neural network with three output heads corresponding to donor, acceptor, and non-splice classes. We additionally evaluated a fine-tuning setting in which all Parnet parameters were updated end-to-end (see Methods). An analogous frozen-versus-fine-tuning strategy was applied to RiNALMo. For Spliceator, which uses separate neural networks for donor and acceptor prediction, we adapted the architecture to a unified binary setting and retrained it on the human subset of the data for fair comparison.

**Figure 5:**
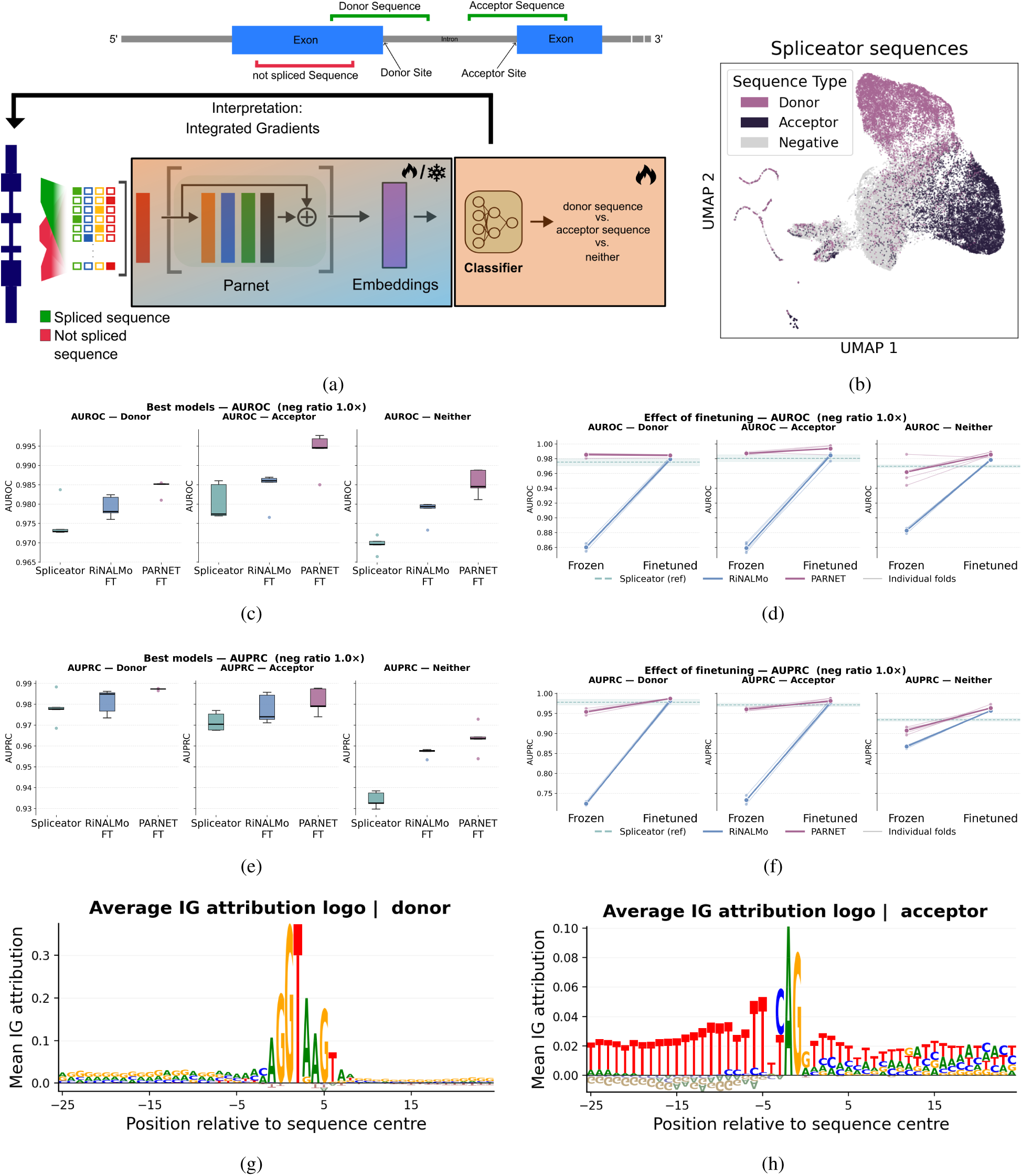
(a) Overview of the downstream ternary classification task distinguishing donor splice sites, acceptor splice sites, and non-splice sites (SS), using either frozen PARNET embeddings or end-to-end fine-tuning of PARNET; (b) Two-dimensional UMAP projection of the PARNET embedding space, where each point represents a sequence and colors denote the true class (donor SS, acceptor SS, or non-SS); (c) Model performance (AUROC) under a 10-fold cross-validation scheme in a balanced setting (equal numbers of positive and negative sequences), comparing Spliceator, Rinalmo, and Parnet using frozen embeddings; (d) Corresponding AUROC results under the same cross-validation setup, comparing Parnet and Rinalmo in both frozen-embedding and end-to-end fine-tuned settings; (e) As in (c), reporting AUPRC instead of AUROC; (f) As in (d), reporting AUPRC instead of AUROC; (g–h) Average sequence logos derived from Integrated Gradients attribution maps for the 500 highest-confidence donor (g) and acceptor (h) sequences.

Parnet frozen embeddings effectively separate sequences around human splice sites from non-splice regions, while also capturing a clear distinction between donor and acceptor sites, as illustrated by the UMAP 2D projection (Fig. 5b). All three benchmarked methods achieve high AUROC and AUPRC on this task (higher than 90%); however, Parnet consistently outperformed the other models across both metrics. Notably, Parnet with frozen embeddings already reaches performance levels that are comparable to, or in most cases indistinguishable from, its fine-tuned counterpart. In contrast, RiNALMo shows a stronger dependence on task-specific fine-tuning, with performance drops of up to 15% in the frozen setting. This highlights the strength of Parnet’s RBP-informed embeddings in capturing biologically meaningful functional structure directly from the pretrained representation space.

### Prediction of intron retention events

After verifying Parnet’s ability to predict donor and acceptor splice sites, we next investigated whether Parnet embeddings capture sequence features associated with alternative splicing events, focusing specifically on intron retention (IR). IR is a form of alternative splicing (AS) in which an intron is not removed from the transcript but instead retained, allowing cells to fine-tune gene expression by reducing transcript stability, preventing export, or modulating translation, often as a way to rapidly regulate protein levels in response to developmental or stress signals [64–66]. Previous studies have shown that IR is regulated not only by the intrinsic strength of the 5*^′^* splice site (SS), but also by a broad range of splice silencing and splice enhancing elements (SSEs and SEEs), which mediate their effects through RBP recognition [67]. One of the major challenges in predicting IR arises from the highly variable lengths of introns, which have been shown to negatively correlate with retention frequency [65, 68, 69]. However, analyses based on artificial splicing constructs have suggested that the majority of splice-regulatory elements are localized within a few hundred nucleotides surrounding the splice sites [70]. Based on these observations, we restricted our analysis to 400 bases on both the 5*^′^* and 3*^′^* ends of each intron, including 100 bases extending into the flanking exons. To integrate information from both intronic boundaries, we concatenated the embeddings derived from the two regions into a joint representation. On top of these Parnet embeddings, we trained a two-layer perceptron (MLP) to classify retained versus spliced introns (see Methods; Figure 6a).

**Figure 6:**
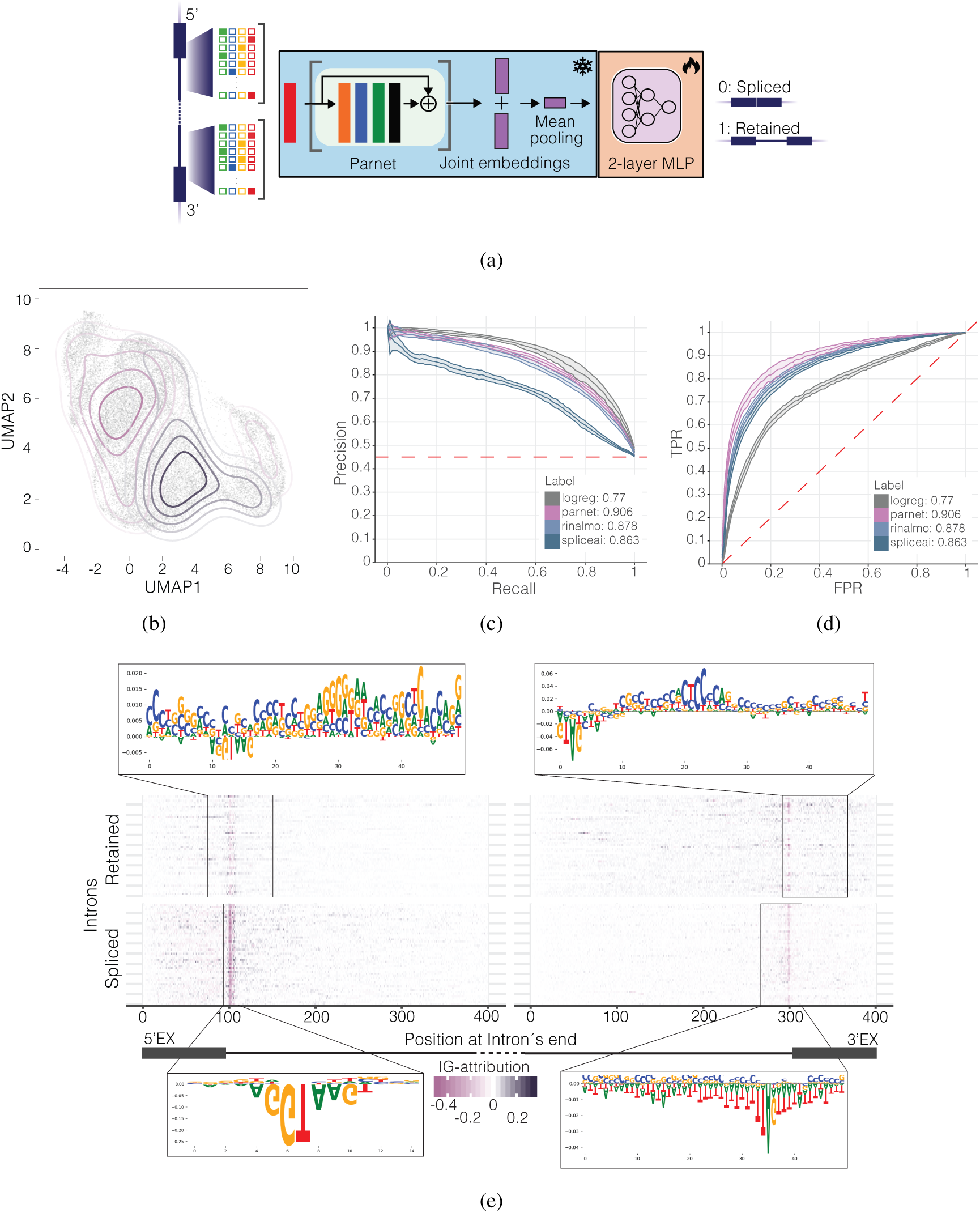
Intron Retention. **a** A schematic illustrating the downstream intron retention prediction task. The hot encoded sequences of the intron’s termini are passed through the Parnet model to generate joint embeddings. The latter are mean-pooled and passed to the 2-layer MLP to predict intron’s binary class (retained vs not retained). **b** Two-dimensional UMAP projection of the Parnet embedding space, where each points represents an intron in the dataset. Contour lines correspond to the two-dimensional densities of each class’ introns. **c** Precision–Recall (PRC) curves across ten folds of zero-shot intron retention (IR) prediction. Different model architectures are indicated by color. Solid lines represent the mean performance across folds, while shaded regions denote the interquartile range (IQR). Top-k performance metrics for each model are reported in the legend. **d** Receiver Operating Characteristic (ROC) curves across ten folds of zero-shot intron retention (IR) prediction. Different model architectures are indicated by color. Solid lines represent the mean performance across folds, while shaded regions denote the interquartile range (IQR). Top-k performance metrics for each model are reported in the legend. **e** Integrated Gradients (IG) attribution maps for PARNET. Per-base attribution scores are shown for the 400 nt flanking each intron end, aggregated across the top 50 predicted introns per class. Positive (retained) and negative (spliced) attributions indicate sequence features driving class predictions. TF-MoDISco motifs extracted_4_f_0_rom the attribution maps are shown above (retained) and below (spliced) the IG maps.

Parnet captures information about retained introns in its embedding space (Figure 6b) and consistently achieves strong classification performance, reaching a top AUROC of 0.90 and a top auPRC of 0.91 (Figure 6c,6d). In the zero-shot setting of distinguishing retained from non-retained introns, we benchmarked against SpliceAI-400 (SpliceAI trained on 400 nt sequences) and RiNalmo. We evaluated both linear classifiers trained on frozen embeddings and downstream MLP heads to assess the additional information encoded in the representations. Both SpliceAI and RiNALMo visually separate the two intron classes in their embedding spaces (Figure **??**,**??**, respectively). While SpliceAI and RiNALMo show broadly comparable performance in the linear setting, with RiNALMo marginally outperforming SpliceAI in the MLP setting (RiNALMo top-k AU-PRC = 0.878; SpliceAI top-k AU-PRC = 0.863), Parnet consistently outperforms both across all evaluation settings (Figure **??**).

We next assessed the sequence determinants of IR learned by Parnet by applying Integrated Gradients (IG) with respect to the target logit [71]. The resulting attribution matrices were then analyzed using TF-MoDISco to identify and map important attribution patterns to specific sequence elements [https://github.com/kundajelab/tfmodisco]. The attribution profiles showed clear signal spikes at both splice sites and correctly identified the canonical splice site sequences as well as a uridine-rich polypyrimidine tract as the main determinants of efficient splicing (6e -bottom part). Conversely, IR was attributed to deviation of the donor site from its canonical AGGUAAGU sequence and higher density of GC-rich elements around the acceptor splice site, together with a weak polypyrimidine tract, in accordance with published reports (Figure 6e - top part) [72], [73], [74]]. Attributions showed presence of both Splicing Silencing Elements and Splicing Enhancing Elements around the 5’ SS in both retained and spliced introns (6d - left part). To contextualize the identified sequence determinants, we mapped TF-MoDISco motifs to the most complete and up-to-date compendium of RBP motifs euPRI with TomTom [54],1 [75]]. Several of the top candidates are established splicing regulators, including hnRNPH1 (binds G-rich elements and modulates splice site choice) and ELAVL2/ELAVL3, which have documented roles in shaping alternative splicing programs [76]. In addition, factors such as RBM15B connect splicing regulation with the m6A writer machinery, providing a plausible mechanistic route by which local sequence context and RNA modification-associated recruitment can bias spliceosome assembly and contribute to IR [77]]. Overall, Parnet enables accurate prediction of intron retention events while remaining interpretable through attribution-based analyses. These analyses recover key regulatory features, including canonical splice-site and polypyrimidine tract signals, as well as GC-rich, splice-proximal RBP-binding motifs, linking predictive gains over other tools to established cis/trans mechanisms of IR regulation.

### Parnet enables zero-shot classification of functional Single Nucleotide Variants (SNVs)

Because Parnet is a sequence-to-signal model, it can predict regulatory signals directly from previously unseen sequences, including reference sequences altered by genetic variants. This capability is particularly valuable for interpreting non-coding mutations, whose functional consequences remain poorly annotated. As shown in our previous analyses, Parnet learns a rich representation of splicing regulatory mechanisms, motivating us to leverage the model in a zero-shot setting to predict the impact of mutations on RNA splicing without requiring variant-specific training. To evaluate Parnet for variant effect prediction, we analyzed the MutSpliceDB dataset, comprising 428 mutations with experimentally validated effects on splicing (236 donor-site and 189 acceptor-site variants). We quantified the impact of each mutation using both changes in Parnet embeddings and predicted RBP binding profiles (see Methods), and compared performance against SpliceAI, RiNALMo, and phyloP across 100 vertebrates. Changes in Parnet embeddings substantially outperformed all competing methods in distinguishing splice-altering MutSpliceDB variants from nearby likely benign variants in GnomAD, achieving an AUPRC of 0.801 and surpassing the current state of the art SpliceAI (Figure 7a). While the aggregated impact on binding profiles shows lesser performance (*auPRC* = 0.68, Figure 7a), it still outperforms conventional PhyloP evolutionary conservation scoring. This suggests that Parnet identified relevant functional signal properties not captured by position-wise evolutionary sequence constraint. Additionally, *in silico mutagenesis* of all 101 positions in vicinity of each of the splice sites affected by MutSpliceDB variants revealed that 358 out of 425 (84.2%) are within the top 10% of most impacting mutations, compared to 150 (6.0%) out of 2,488 for the GnomAD controls. This further confirms the disruptive nature of the MutspliceDB variants.

**Figure 7:**
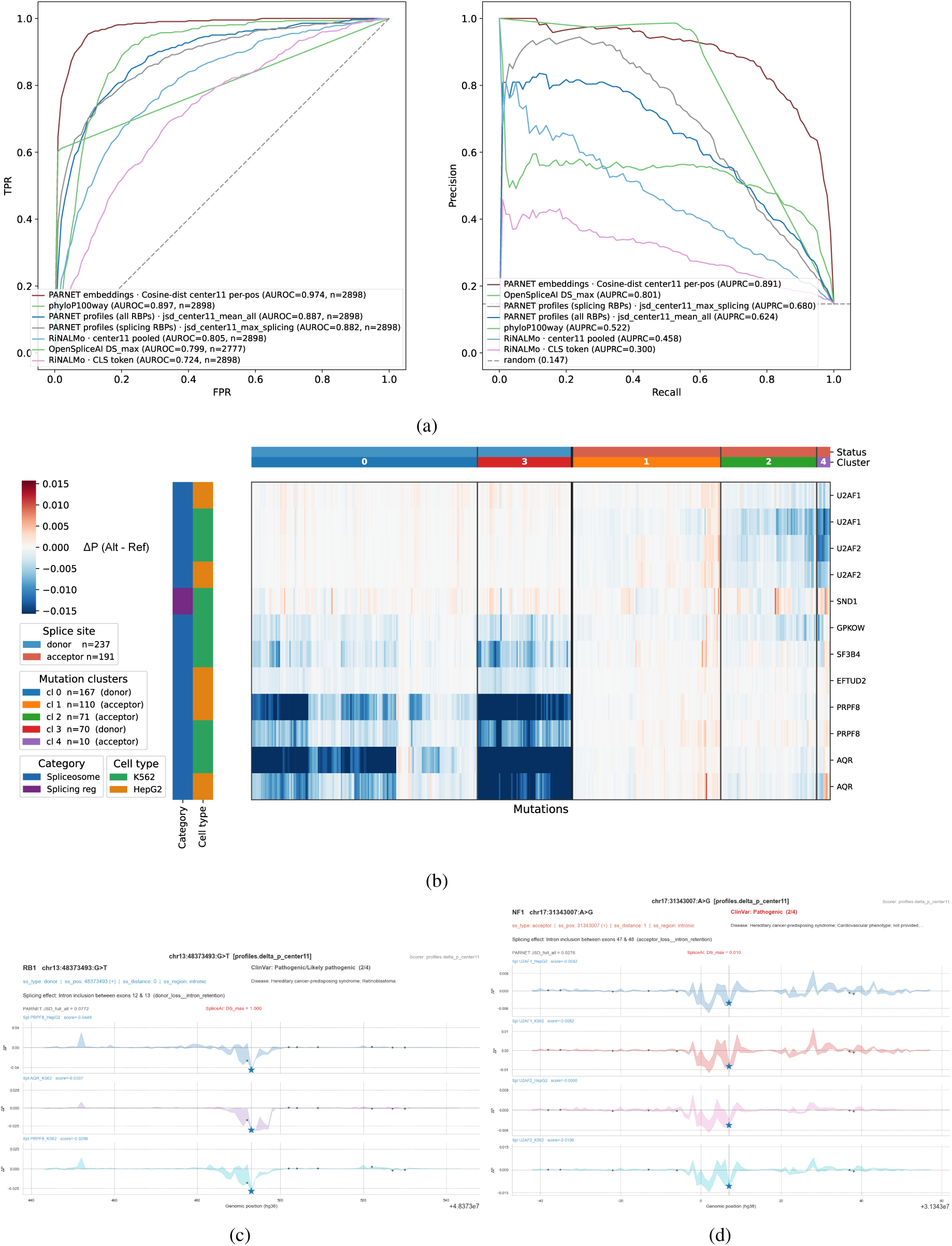
Classification of splicing variants with the PARNET model. Figure 7**. A** auROC and auPRC for evaluating the performance of different scoring approaches to classifying experimentally validated splicing-impacting mutations against local GnomAD SNVs. Cosine similarity between REF and ALT sequences was used to score mutations, using the embeddings representations from PARNET, RinALMo. Additionally, functional scores are reported from PARNET’s RBP binding profiles, as well as from SpliceAI mutation scores. Sequence Conservation scores as measured by PhyloP are also included for comparison. **B** Heatmap showing the impact of MutSpliceDB mutations (rows) on the 12 most affected RBP binding profiles (columns), quantified as the change in PARNET-predicted binding score within an 11-nt window centered on each mutation. Mutations are grouped by donor or acceptor splice-site status, with cluster annotations indicated by the color bar above the heatmap. RBPs are annotated by functional class (spliceosome or splicing regulator) and by the cell line from which the corresponding eCLIP data were generated (color bars on the left side). **C, D** Examples of binding impact in the local *±*) 50 nucleotide around two selected mutations. The binding impact of all possible mutations in those regions was evaluated through *in silico* mutagenesis, computing for each mutation the difference in cumulated binding probability over a 11 nucleotide interval centered on the mutation. The splicing mutations are located at the center, while local GnomAD controls are shown for reference (grey points). (Left) A donor site mutation is associated with decrease in binding of PRPF8 and AQR. (Right) An acceptor site mutation is associated with decrease in binding of 3’site definition RBPs.

Beyond predicting the functional impact of a variant, Parnet directly estimates how the mutation alters RNA-binding protein (RBP) binding, providing mechanistic insight into the regulatory processes disrupted by the variant. As illustrated in Figure 7b, hierarchical clustering of MutSpliceDB mutations based on predicted changes in RNA-binding protein (RBP) binding profiles (restricted to the top 12 RBPs) reveals a clear organization of variants into a small number of distinct functional groups. For donor-site mutations, two main clusters can be distinguished, corresponding to high (cluster 3) and moderate (cluster 0) disruption of core spliceosomal RBPs, including AQR and PRPF8 [78], consistent with graded effects on spliceosome assembly. Acceptor-site mutations similarly separate into three clusters: two exhibit strong and moderate decreases in binding of the canonical 3*^′^* splice-site recognition factors U2AF1 and U2AF2 [79], while a third cluster comprises mutations with weaker or more heterogeneous effects, which are more difficult to clearly associate with canonical RBP disruption patterns.

To further show the value of Parnet for interpreting mutation effects on RBP binding, we selected two clinically relevant splice-site variants and identified the RBPs predicted to be most strongly affected. The first variant is a donor-site substitution in RB1 (NM_000321.3:c.1215+1G*>*T, Fig. 7c). Its reported effect in MutspliceDB [80] is the retention of the intron between exons 12 and 13, as measured from lung tumour RNA-seq. Two independent clinical laboratories report it in germline testing for retinoblastoma and hereditary cancer predisposition, and ClinVar [81] aggregates it as pathogenic/likely pathogenic (ClinVar VCV001298329). Through *in silico mutagenesis* (ISM) we confirm that the mutation has the largest impact compared to any other mutated position in vicinity, as measured for PRPF8 and AQR. The second variant is an acceptor-site substitution in NF1 (NM_001042492.3:c.7063-2A*>*G, Fig. 7d). Its reported effect is the retention of the intron between exons 47 and 48. Besides, it is classified as pathogenic in germline testing for neurofibromatosis type 1 and hereditary cancer predisposition (ClinVar VCV000635820). Interestingly, this mutation is missed by SpliceAI, which scores it as very low impact. The *in silico mutagenesis* reveals that the mutation lies within a functionally variable region where mutations are predicted to have either enhancing or decreasing binding effect on the U2AF2 and U2AF1 RBPs. While the mutation itself is predicted to decrease the binding of these two RBPs at the location of the mutation, the overall ISM profiles highlight that 3*^′^* splice sites represent a more diffuse functional region, from the branch point to the polypyrimidine tract and the AG di-nucleotide itself.

We extended our analysis of mutation impact to the datasets gathered in the SpliceBench resource [82]. Briefly, this resource gathers variants and their functional status from four Minigene Splicing Reporter Assays MSRA) studies as well as one Saturation Genome Editing study of the BRCA1 gene and one manually-curated set of variants in the MLH1 gene. On the aggregated dataset, Parnet ranks second (8a, AUROC=0.782, AUPRC=0.661, discarding CADD which is only available for BRCA1 mutations), outperformed by SpliceAI (AUROC=0.794, AUPRC=0.709). Across all dataset, Parnet ranks in the top 3 (as per AUPRC) in 4 out of 6 datasets.

Similarly to the MutspliceDB mutations, we explored how the individual RBPs were impacted by mutations, identifying for each dataset the most discriminative tracks. As seen on the RON (or MST1R) exon 11 (8h), the Parnet cosine distance calculated on embeddings of reference and mutated sequences highlights particularly well the boundaries of the exon 11 (and outperforms all other methods), while top RBPs include the spliceosomal factors AQR and SMNDC1, as well as GEMIN5, a factor involved in snRNP biogenesis. Similarly, exploring the profiles of impact over the exon 9 of WT1 (8h) shows high putative impact measured by Parnet nearby the exon ends, with predicted decreased binding for AQR and PRPF8 at those locations.

Overall, we demonstrate that Parnet can effectively discriminate functional mutations in a zero-shot setting and accurately identify the RBPs most strongly impacted by each variant. At the same time, interpretation of RBP-specific effects should be approached with care, as eCLIP-seq data may reflect not only direct binding specificity but also transient physical proximity of RBPs to RNA elements involved in splicing. Additional factors such as splicing kinetics and differences in cross-linking efficiency may further influence these signals. As a result, Parnet may capture a mixture of sequence-driven proximity effects rather than purely mechanistic binding preferences. Nevertheless, despite these caveats, the strong discriminative performance of the model indicates that its predictions of overall perturbations to the splicing machinery robustly capture the functional consequences of mutations.

## Discussion

In this work, we introduced Parnet, a multi-task foundation model for RNA representation learning trained end-to-end on 223 eCLIP-seq experiments to predict base-resolution RBP binding profiles from raw RNA sequences. The main contributions of this work are: (i) a novel pretraining strategy that grounds RNA sequence representations directly in experimentally measured protein–RNA interactions, providing stronger biologically motivated inductive biases than self-supervised reconstruction objectives; (i) A multi-task extension of RBPNet that substantially improves both binding profile prediction accuracy and RBP sequence motif recovery compared with single-task RBP models, while also enabling the prediction of functional RNA-binding sites associated with RNA regulatory events, such as regulated exon retention and exon skipping; (ii) a novel pretraining strategy that grounds RNA sequence representations directly in experimentally measured protein–RNA interactions, providing stronger biologically motivated inductive biases than self-supervised reconstruction objectives; (iii) demonstration that Parnet embeddings, encoding the combinatorial “RBP code” governing post-transcriptional regulation, can be used to reach state-of-the-art or superior performance across a variety of prediction tasks with minimal or no fine-tuning, notably outperforming much larger self-supervised RNA and genomic language models; and (iv) a mechanistic interpretability framework in which predictions of RNA processed can be traced back to specific RBPs and their sequence motifs via integrated gradients and embedding space interrogation, directly linking computational predictions to the molecular machinery governing post-transcriptional regulation.

### Multi-task learning improves modeling of eCLIP signal

Parnet extends RBPNet from single-task RBP binding prediction to joint modeling of 223 eCLIP experiments. This multi-task formulation improves performance on held-out transcripts, increasing average Pearson and Spearman correlations by *∼*35% and *∼*19%, respectively, while also improving peak and cross-link identification. The largest gains are observed for the lowest-quality datasets: RBPs in the bottom performance quartile improve by *∼*40% in Spearman correlation, compared with only *∼*3.6% for the top quartile. This trend suggests that parameter sharing enables poorly sampled experiments to leverage information from related RBPs with similar binding preferences, consistent with the association between performance gains and data-quality proxies (peak count, unique-read fraction). However, Parnet also has substantially greater capacity (*∼*21M vs. *∼*0.5M parameters) and a larger context window, so the improvements cannot be attributed to multi-task learning alone.

Beyond predictive accuracy, the *λ* penalty on target–control mixing provides an orthogonal, mechanistically interpretable quality check. As *λ* increases, the target track sheds shared background signal (e.g. the G-rich pattern for RBFOX2, U-rich for QKI) into the control track, and denoised 5-mer preferences show progressively stronger concordance with in vitro motifs (RNA Bind-n-Seq, RNAcompete) up to *λ* = 10, beyond which excessive shrinkage degrades the signal. This convergence toward assay-independent binding specificities is a strong evidence that the learned representations capture intrinsic sequence recognition, not experiment-specific artifacts. Robustness across cellular contexts and CLIP protocols is exemplified by TDP-43 binding at the STMN2 cryptic exon: although STMN2 is expressed in neither the K562 nor the HepG2 cell lines of the eCLIP data used for training, Parnet’s predicted profile recovers the TDP-43 iCLIP signal measured in SH-SY5Y neuroblastoma cells, transferring across both assay type and cell context without retraining.

Predicted profiles also recover the regulatory grammar of splicing. RNA maps built from Parnet predictions reproduce the position-dependent logic of diverse regulators, from 3SS enhancement by SF3B4 and U2AF2 to exonic enhancement by the SR proteins TRA2A and SRSF7, 5SS enhancement by TIA1 and RBM22, and 3SS silencing by PTBP1 and HNRNPK. Signal rises at regulated exons without inflating the false-positive rate, and because the model places binding relative to each splice site rather than merely detecting it, these profiles reflect functionally coherent recognition rather than aggregated occupancy.

Overall, Parnet not only predicts binding profiles more accurately than single-task models, but learns representations that recover intrinsic sequence specificity and transfer to unseen assays, cell types, and regulatory contexts.

### Functional pretraining as an alternative to self-supervised reconstruction

The dominant paradigm for RNA foundation models has been masked language modeling (MLM) on large unlabeled sequence corpora, as exemplified by RNA-FM [41], RiNALMo [42], ERNIE-RNA [43], or Uni-RNA [83]. These models learn representations that capture sequence conservation and structural context, achieving strong performance on RNA structure prediction and ncRNA classification. However, MLM objectives are agnostic to biological function, treating all nucleotide positions equally despite the fact that only a subset directly contributes to RNA regulation. This likely limits the functional specificity of the learned representations and explains why these models often require extensive fine-tuning and large labeled datasets for downstream regulatory tasks. For example, in splice site prediction, RiNALMo requires substantial fine-tuning to match the performance of Parnet.

Parnet represents a different class of foundation model. Instead of a function-agnostic objective, it learns its representation from an experimental measurement of function, RBP binding, pretraining in a supervised manner exclusively on RBP-bound RNA sequences. This grounds the model in RNA processing and regulation directly, rather than leaving functional structure to be recovered during fine-tuning. As a result, it learns functionally informative representations that generalize across diverse RNA regulatory tasks without large-scale fine-tuning. Consistent with this, frozen Parnet embeddings match or outperform other RNA language models, including on tasks outside its pretraining objective. Embedding analyses further support this interpretation: removing the top principal components substantially reduces the performance of RiNALMo and Nucleotide Transformer, indicating reliance on global sequence features such as length and GC content, whereas Parnet is largely unaffected, suggesting it encodes more functionally specific information.

### The RBP interactome as a sufficient basis for functional RNA representations

A central hypothesis underlying Parnet is that combinatorial RBP binding provides a highly informative proxy for RNA function, as RBP occupancy governs splicing, stability, translation, and other key regulatory processes. Our results provide strong evidence in support of this hypothesis. Parnet, trained exclusively on RBP binding data from two cell lines and 223 eCLIP experiments, generalizes to predict intron retention events, translational efficiency, and splice site sequences from different cell types. Notably, Parnet enables zero-shot prediction of lncRNA chromatin dissociation half-lives, separating chromatin-retained from nucleoplasmic lncRNAs without fine-tuning while recapitulating key determinants identified by chrTT-seq, including SNRNP70 enrichment in retained transcripts and SRSF factors in rapidly released ones. This demonstrates that Parnet has internalized higher-order regulatory logic that was never explicitly part of its training objective. Mechanistically, the model recapitulates position-dependent splicing principles, including distinct binding patterns of SR proteins, hnRNPs, spliceosomal components, and TIA1 around regulated exons, demonstrating that its representations encode genuine biological knowledge rather than generic sequence features.

### Interpretability through the RBP lens

A critical distinctive feature of Parnet is that its interpretability is mechanistically grounded in a way that is unavailable to self-supervised models. Because Parnet is trained to predict RBP binding, its embedding space can be directly interrogated by projecting sequences and overlaying RBP occupancy scores, yielding intuitive associations between embedding neighborhoods and specific regulatory proteins. In the translational efficiency tasks, this approach identified LARP4, EIF3H, and EIF4G2 as positively associated with high-TE transcripts, all well-established components of the translation initiation and ribosome recruitment machinery, while RBM15 enrichment in low-TE regions is consistent with its role in m6A-mediated modulation of RNA stability. These associations were recovered purely from the Parnet embedding geometry, without any direct supervision on translation. In the IR task, Integrated Gradients followed by TF-MoDISco and TomTom matching recovered canonical splice site sequences, polypyrimidine tracts, G-rich elements associated with hnRNPH1, and RBM15B as a link between splicing regulation and the m6A writer machinery, all mechanistically plausible and in agreement with published literature. This capacity to trace predictions back to specific RBPs and their binding motifs represents an interpretability advantage over current RNA foundation models.

### Comparison with philosophically related models

Parnet is conceptually most closely related to two lines of work that have pursued functionally motivated alternatives to self-supervised genomic pretraining. Borzoi and BigRNA ground DNA-level foundation models in experimental functional genomics tracks, such as chromatin accessibility, transcription factor binding, and gene expression, achieving foundation-like versatility while remaining anchored to measured biology. Parnet extends this philosophy into the post-transcriptional domain, replacing chromatin-level assays with CLIP-seq data as the functional pretraining signal. Among RNA foundation models, Orthrus offers an instructive contrast to Parnet. Instead of masked language modeling, Orthrus uses contrastive learning on mRNA isoforms and evolutionarily related transcripts from over 400 mammalian species, leveraging conservation as a proxy for function. In contrast, Parnet learns functional representations directly from experimentally measured protein–RNA interactions and provides direct mechanistic interpretability at the level of individual RNA-binding proteins.

### Limitations and scope

Despite its strong performance, Parnet has some limitations that should be acknowledged. First, the training data covers only 150 RBPs across two cell lines (K562, HepG2) for a total of 223 eCLIP experiments, representing approximately 10% of the estimated RBP repertoire. The remaining 90% of RBPs, for which no CLIP data currently exists, are absent from the training signal. It is therefore possible that regulatory programs governed predominantly by uncharacterized RBPs are underrepresented in Parnet’s embedding space. However, the consistently strong downstream performance we observe may suggest a degree of redundancy in the RBP code: the sequence features recognized by the 150 profiled RBPs may be sufficient to implicitly encode the regulatory logic of a much larger fraction of the transcriptome. Nevertheless, interpretation is inherently biased toward the profiled RBPs, and caution is needed when using Parnet attributions to draw mechanistic conclusions in biological contexts that differ substantially from ENCODE K562 and HepG2 cell lines. Second, Parnet currently operates on sequence alone and does not incorporate RNA secondary structure or chemical modifications such as m6A. These features that are known to modulate RBP binding and RNA fate *in vivo*. The recurrent association of m6A-related RBPs (RBM15, RBM15B) with functional predictions hints that Parnet partially captures modification-associated regulatory logic indirectly, through the binding footprints of modification-associated RBPs, but explicit modeling of these modalities could further enrich the representations.

### Future directions

Several natural extensions follow from the current work. The most impactful would be to expand the RBP coverage beyond those with available CLIP data by learning joint protein–RNA representations. By simultaneously embedding RBP sequences or structures alongside RNA sequences in a shared latent space, it would be possible to infer binding preferences for uncharacterized RBPs from their sequence homology to profiled ones, dramatically broadening the functional scope of the training signal. Such a multimodal architecture would also create a natural framework for RNA sequence design: because Parnet simultaneously models binding profiles for hundreds of RBPs, Several natural extensions emerge from this work. First, Parnet could be expanded beyond RBPs with available CLIP data by learning joint protein–RNA representations, enabling prediction of binding preferences for uncharacterized RBPs from sequence or structural similarity. Such a multimodal framework would also support RNA sequence design by optimizing RBP occupancy profiles to engineer regulatory properties such as splicing, stability, and localization. The intron retention results suggest an immediately practical application: by examining which RBPs drive Parnet’s predictions for any set of differentially retained introns or exons of interest, researchers can generate experimentally testable hypotheses about the regulators responsible for the observed splicing changes, a generalization of the RNA map concept to arbitrary RNA processing events. Finally, integrating RNA structure and modification information as additional input modalities or auxiliary pretraining objectives offers a natural path toward more comprehensive *in vivo* RNA representations.

## Conclusion

In this work we develop Parnet, a CLIP-seq–driven RNA foundation model that learns functional representations of RNA sequences by predicting the binding profiles of hundreds of RBPs simultaneously. Parnet demonstrates that anchoring foundation model training directly in experimental functional data, such as the the RBP interactome, is a powerful and efficient alternative to self-supervised sequence reconstruction for learning RNA representations. With 21M parameters and training data limited to two cell lines, Parnet reaches state-of-the-art performance across a diverse range of RNA functional prediction tasks, outperforming other RNa language models that are substantially larger and trained on orders of magnitude more sequence data or task-specific models. Its representations are inherently interpretable through the lens of protein–RNA interaction biology. We anticipate that Parnet will serve as a versatile foundation for computational RNA biology, from the prediction and interpretation of post-transcriptional regulatory events to the prioritization of disease-associated non-coding variants and the design of RNAs with tailored regulatory properties.

## Methods

### Data Pre-processing

A total of 223 eCLIP datasets were obtained from the ENCODE [84] database, encompassing 150 unique RBPs (103 and 120 for HepG2 and K562 cell lines, respectively). eCLIP data was processes as described in [46]. In brief, for each eCLIP experiment and size-matched input (SMI) control, BAM files of both biological replicates were merged. Subsequently, aligned R2 reads where extracted and 5’ read-starts were counted at each nucleotide position in a stranded manner.

To construct samples for model training and validation, coding and non-coding transcripts were extracted from GENCODE (*v48*) and overlapping annotations were merged. The resulting regions were then evenly tiled into 600*nt* long sequences with an overhang of 150*nt* for adjacent tiles. Furthermore, a minimum filter of 8 read-counts over the length from any track (eCLIP or SMI) of any experiment was applied so as to retain tiles with available signal to learn from. This resulted in a total of 700,114 tiles. All sequences were padded to a length of 600 using *N* (unknown nucleotide) symbols. Padding was applied at the 5*^′^*, 3*^′^* or both sides, depending on the location of each tile within the region.

Tiles were then split into train, validation and test sets, using the same chromosome-based splitting strategy described in [46], i.e. isolating sequences from chromosomes 3, 8, and chr15 for the test set, sequences from chromosomes 2, 9 and 16 for validation, and sequences from other autosomes for training set.

### Parnet Model Architecture

The Parnet architecture builds on the model backbone of RBPNet [46] with significant changes. We extended the length of RNA sequences for training from 300 to 600 to account for a larger context, and up-scaled the number of convolutional filters and residual blocks in order to account for the significantly larger number of experimental track the models has to predict. As a result, the model size increased from 0.5 Million to 21 Million parameters. Briefly, RNA sequences are one-hot encoded and processed by an initial 1D convolutional layer (128 filters, kernel size 12), followed by nine residual blocks. Each block comprises a dilated 1D convolution (128 filters, kernel size 6, exponentially increasing dilation), batch normalization, ReLU activation, and 25% dropout. From the bottleneck layer of the model’s backbone, the model branches into 223 output heads, one for each experiment. Each output head consists of a transposed 1D convolution with a single filter (kernel size 20), producing a 600-position probability distribution over the input sequence.

Because experimental biases generate nonspecific eCLIP signal, matched control experiments are used to estimate locus-specific background. Similar to RBPNet, each head in Parnet consists of a control track, modeling the crosslink count distribution of the SMI experiment, a target track, which represents the unbiased, protein-specific distribution, and the total track, which models the eCLIP crosslink counts. The total track is obtained through additive mixing of the control and target tracks using a mixing coefficient *π*, i.e., *p_total_* = *π × p_target_* + (1 *− π*) *× p_control_*, where *p_total_* is the probability distribution of the observed total (e.g., eCLIP) signal, and *p_control_* and *p_target_* are the probability distributions for the control and latent target signals, respectively, learned directly from sequence.

### Mixing Coefficient Penalty

The goal of the above formulation is to disentangle the crosslink count signal into protein-specific binding and experimental bias components. We assume that for each sample there is some mixing coefficient *π^′^* at which the model makes maximal use of the control track to explain experimental biases in the eCLIP track, such that the target track purely contains protein-specific binding information. In practice, the mixing coefficient is unconstrained and information from the SMI track may leak freely into the target track, such that the actual mixing coefficient may settle between *p^′^* (optimal use of control track) and 1 (no use of control track, *p_total_*= *p_target_*). In order to rectify this and steer *π* towards *π^′^*, we introduce an additional penalty term, *λπ*, to the model’s loss formulation, where *λ* controls the strength of this penalty. By by using the magnitude of the mixing coefficient as an additional loss term, the model is incentivized to make maximal use of the control track in order to keep the loss (and therefore the mixing coefficient) small.

### Multi-task Loss

Following RBPNet, losses for the eCLIP and SMI tracks are defined as the negative log-likelihood *−* log *p_mult._*(*c_obs_ | p_pred_, n_obs_*), for each experiment and track. Together, the final model loss is given by

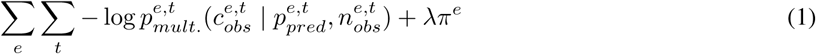

for each experiment *e* and track *t ∈ {eCLIP, SMI}*.

### Model Training

All models were trained using the AdamW optimizer [85] with a learning rate of 0.001 for at most 100 epochs. Training was terminated if the model did not improve for the last 10 epochs and the best-performing model on the validation set was saved as the final model.

### Model evaluation

#### Performance against eCLIP signals

Performance of the model was evaluated by correlating the predicted binding profiles (total tracks) with read start counts (summed over replicates or from each separated replicate) for each 600-nt held-out test sequence, and averaging PCCs across all sequences for each eCLIP experiment. We report both Spearman and Pearson correlations.

In addition, we quantified the enrichment of the predicted probability density over annotated binding intervals. Specifically, we treated the probability density as a continuous scoring function and evaluated its ability to distinguish binding from non-binding positions using Receiver Operating Characteristic (ROC) and Precision–Recall (PR) curves. As ground truth labels, NarrowPeaks from the ENCODE database were gathered and merged from the two replicates as a first set of ground truth binding intervals. To better capture the single-nucleotide resolution of Parnet predictions, we repeated this analysis using single-nucleotide crosslink (CL) sites identified by the PureCLIP software with default parameters ([48]). Each called CL site was extended by 5 nucleotides on either side to define the ground truth regions.

#### Performance against peak and single-nucleotide binding site annotations

Performance of the model was also evaluated by comparing predicted probabilities with ENCODE NarrowPeaks (called by CLIPper [47] and with crosslink (CL) sites identified by PureCLIP [48].

To this end, we performed genome-wide peak calling with PureCLIP on ENCODE eCLIP datasets. We intersected held-out chromosome transcripts with either ENCODE peaks or PureCLIP CL sites. Parnet was then used to predict transcript-wide binding profiles. For each sequence interval, we computed the area under the Receiver Operating Characteristic curve (AUROC) and average precision (AP) by treating positions overlapping ENCODE peaks or PureCLIP CL sites as positives and all others as negatives, considering these annotations as ground truth. AUROC and AP were computed per sequence and then averaged to obtain RBP-specific performance metrics.

### Downstream Tasks

#### Genomic regions and RNA family classification

**Datasets** Sequences were extracted from the human genome (GRCh38). Nucleotide sequences and annotations were downloaded from GENCODEv40 ([86]). For the RNA family classification task, we selected five biotypes: protein-coding transcripts, long non-coding RNAs (lncRNAs), pseudogenes, microRNAs (miRNAs), and a non-coding RNA class (ncRNA). This last class was further subdivided into small nucleolar RNAs (snoRNAs), small nuclear RNAs (snRNAs), small cajal body-specific RNAs (scaRNA), ribosomal RNA (rRNA), mitochondrial transfer RNA (Mt_tRNA), ribozyme, small RNA (sRNA), mitochondrial ribosomal RNA (Mt_rRNA), miscellaneous RNA (miscRNA) and others. For the genomic region classification task, sequences were annotated as coding sequences (CDS), 5‘ untranslated regions (5‘ UTR), 3‘ untranslated regions (3‘ UTR), and introns.

**Task description** To evaluate the biological information captured by Parnet embeddings, we performed classification tasks on RNA families and genomic regions. Each nucleotide sequence was one-hot encoded and passed through the frozen Parnet encoder. Sequence-level representations were obtained by mean pooling the nucleotide-wise embeddings across the full sequence length. To assess the contribution of confounding factors to the embedding space, we computed Pearson correlations between the first four principal components (PCA) of the embeddings and two sequence-level covariates: GC content and transcript length. We additionally trained classifiers on embeddings with the first two principal components removed (*PCA-50 Skip 2*) to evaluate classification performance after decoupling from these confounders. Classification was performed using a two-hidden-layer neural network with layer sizes 256 and 128, followed by a softmax output layer. Sequence embeddings were classified using multinomial logistic regression. The dataset was partitioned into a training set (80%) and a held-out test set (20%) using a stratified split to preserve class proportions. Model performance was first assessed on the training set via stratified 10-fold cross-validation. A final classifier was then refit on the complete training set and evaluated on the independent test set. Classification performance was quantified per class using the area under the receiver operating characteristic curve (AUROC) and the area under the precision-recall curve (AUPRC) in a one-vs-rest scheme, in which each class in turn was treated as the positive class and all remaining classes as the negative class. Overall performance was additionally summarized by accuracy and class-weighted precision, recall, and F1-score.

**Comparison with other models** We benchmarked Parnet against two pretrained nucleotide language models: Ri-NALMo [42], giga-v1 and the Nucleotide Transformer [38] 500M_human_ref. For both baselines, we extracted nucleotide-level embeddings and used them as fixed input representations for the downstream task, replacing Parnet’s learned representations while keeping the rest of the evaluation pipeline identical. Sequences shorter than each model’s maximum context length were processed in a single forward pass, and the resulting per-token hidden states were used directly as the sequence representation. For longer sequences, we split the input into non-overlapping chunks corresponding to each model’s maximum context: 1, 024 nucleotides for RiNALMo and 5, 994 nucleotides for the Nucleotide Transformer. Each chunk was processed independently, and the resulting per-chunk embeddings were mean-pooled position-wise across chunks to produce a fixed-length representation matching a single chunk’s output. To ensure a fair comparison, the same two-hidden-layer classifier (layer sizes 256 and 128, followed by a softmax output layer) was applied on top of the embeddings from Parnet, RiNALMo, and the Nucleotide Transformer. Model performance was evaluated using the area under the receiver operating characteristic curve (AUROC), computed per class.

#### Prediction of lncRNA Localization Dynamics

**Datasets** Long non-coding RNA localization dynamics data, under the form of chromatin dissociation kinetics, along with the lncRNA sequences were taken from Ntini et al.[56]. In that study, nascent RNA dissociation from chromatin, i.e. chromatin dissociation half-life, was measured genome-wide in MCF7 cells using chrTT-seq, a pulse-chase metabolic labeling approach that tracks the ratio of chromatin-associated to total nuclear RNA over time. Per-transcript chromatin dissociation half-lives (*t*_1_*_/_*_2_) were pre-computed by fitting an exponential decay model

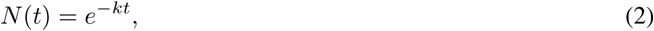

to the time-series data, where *k* is the dissociation rate and *t*_1_*_/_*_2_ = ln(2)*/k*. Transcripts with slow dissociation kinetics were defined as ‘chromatin-retained’, while those with fast kinetics as ‘nucleoplasmic’.

**Task description** Each lncRNA sequence was one-hot encoded and passed through the frozen Parnet encoder, and nucleotide-wise embeddings were mean pooled over the full transcript length to obtain a fixed-length sequence representation. Regression of the pre-computed half-life values was performed using either a linear regressor or a two-hidden-layer neural network with layer sizes of 256 and 128 units followed by a linear output layer, both trained on frozen embeddings only. All regressors were trained and evaluated using 10-fold cross-validation with early stopping, and performance was assessed using Pearson correlation between predicted and observed half-life values. Parnet was compared against Nucleotide Transformer and RiNALMo embeddings under the same setup, with all three models applied to full transcript sequences.

**Comparison with other models** For this task, RiNALMo and the Nucleotide Transformer were used as described in the paragraph above (Section **??**). On top of the resulting embeddings, the same two-hidden-layer neural network (layer sizes 256 and 128) was applied, but with a regression head replacing the softmax output. Model performance was evaluated using Pearson correlation over 10 cross validation folds.

#### Prediction of mRNA Translation Efficiency

**Datasets** From Parnet embeddings, we predicted translational efficiency (TE) across four ribosome profiling datasets spanning different cell types: HEK293T cells from Aleksaki et al.[59] and [87], as well as skeletal muscle and PC3 prostate cancer cells. The Aleksaki dataset was preprocessed by Schlusser et al. [61], while the remaining three dataset were preprocessed by Cao et al.[57]. Per-transcript translation efficiency was computed from matched RNA-seq and Ribo-seq data as *TE* = *RPKM_Ribo−seq_/RPKM_RNA−seq_*, with transcripts lacking sufficient coverage in either assay discarded to avoid unstable ratios. The prediction target was log-transformed translational efficiency (log TE) for all four datasets, defined as defined the amount of ribosome occupancy (or protein synthesis) per unit of mRNA abundance. Transcripts were not matched across datasets; each dataset was treated independently.

**Task description** For all datasets, each transcript sequence was one-hot encoded and passed through the frozen Parnet encoder. Sequence-level representations were obtained by mean pooling over the nucleotide-wise embeddings across the full transcript length, encompassing the 5‘ UTR, CDS, and 3‘ UTR. This is in contrast to baseline models (Optimus 5‘ [60], UTR-LM [62], Framepool [16], and TranslateLSTM [61]), which were trained and evaluated on 5‘ UTR sequences only. The use of mean pooling over variable-length sequences in the CNN-based Parnet architecture naturally accommodates full-length transcripts without requiring fixed-length inputs. Regression was performed using a two-layer neural network with layer sizes of 256 and 128 units, followed by a linear output layer. All regressors were trained with early stopping and evaluated using 10-fold cross-validation. Performance was assessed using Pearson correlation between predicted and observed TE values.

**Comparison with other models** For a fair comparison with Parnet under an identical input regime, RiNALMo [42] embeddings were also computed on full transcripts rather than on the 5‘ UTR alone. For each transcript, the full-length sequence spanning the 5‘ UTR, CDS, and 3‘ UTR was tokenised at single-nucleotide resolution and passed through the frozen RiNALMo encoder. Transcripts exceeding RiNALMo’s maximum context length were split into non-overlapping windows, whose embeddings were averaged to obtain a single transcript representation. These embeddings were then used as input to the same two-layer regression network (256 and 128 hidden units) and evaluated using the identical 10-fold cross-validation and Pearson correlation protocol as Parnet. In contrast, the other four baseline models operate on the 5‘ UTR rather than the full transcript, but they differ in how 5‘ UTR length is handled. Optimus 5‘ and UTR-LM require fixed-length inputs: for both we used the 100-nucleotide variant, so 5‘ UTRs longer than 100 nt were truncated to the 100 nucleotides immediately upstream of the start codon, while shorter 5‘ UTRs were padded at their 5‘ end to the required input length, following the original Optimus 5-Prime convention. In contrast, FramePool and TranslateLSTM natively accommodate variable-length sequences—FramePool through frame-wise pooling and TranslateLSTM through its recurrent architecture—and were therefore applied to full-length 5 UTRs without truncation or padding.

#### Splicing Site Prediction

**Datasets** To evaluate the ability of Parnet embeddings to capture splice-site signals, we used the Spliceator dataset [63], which comprises experimentally validated donor and acceptor splice-site sequences from multiple species. Positive sequences corresponding to the splice sites (SS) were obtained from the G3PO+ benchmark data set, including 10, 000 donor and 10, 000 acceptor sites in humans. We selected the version of the dataset in which each sequence is 400 nucleotides long and centered on the splice site. Negative sequences, as described in the Spliceator study, were randomly sampled from exonic, intronic, or false-positive splice-site regions in G3PO+, centered on GT or AG di-nucleotides that do not correspond to annotated splice sites in the positive set [42].

**Task description** We trained a three-way classifier to simultaneously distinguish donor splice sites, acceptor splice sites, and non-splice sequences (negative class). Each 400-nt sequence was one-hot encoded and processed through the Parnet encoder, yielding nucleotide-wise embeddings of dimension 768, consistent with the fixed-length sequence setup used during training. These embeddings were then used as input to a two-hidden-layer neural network classifier with 256 and 128 units, respectively, followed by a three-class softmax output layer. We evaluated two training regimes:

i. head-only training, in which the Parnet encoder weights were frozen and only the classification head was trained, and
ii. full fine-tuning, in which all model parameters were optimized end-to-end. To assess robustness to class imbalance, models were trained and evaluated using three negative-to-positive ratios: 0.5, 1, and 2, corresponding to half, equal, and twice as many negative samples as positives per splice-site class. All models were trained with early stopping and evaluated using 10-fold cross-validation. Performance was assessed using class-wise AUROC and AUPRC metrics.

**Comparison with other models** For this task, Parnet was compared against RiNALMo and the Spliceator baseline. For Parnet and RiNALMo, the same three-way classification head was used: a two-hidden-layer neural network (layer sizes 256 and 128) followed by a softmax output over the three classes. To ensure a fair comparison with Parnet, RiNALMo was evaluated in two settings: a frozen setting, in which its weights were kept fixed and only the classification head was trained, and a fine-tuned setting, in which its weights were updated jointly with the head during training. For Spliceator, we used the original model architecture and adapted its output layer from binary to three-way classification, training the model from scratch on the data. All models received the same input sequences. Model performance was evaluated using the area under the receiver operating characteristic curve (AUROC) and the area under the precision-recall curve (AUPRC), both computed per class.

#### Intron retention prediction

**Datasets** The consensus retained intron dataset was built from the vast corpus of reprocessed cancer cell line encyclopedia (CCLE) splicing data [88]. An intron was labeled “Retained” if its percent intron retained value (PIR), corresponding to the fraction of transcripts from a gene that retain a specific intron, was *PIR ≥* 30 in *≥* 10 different cell lines. A “Spliced” intron label was assigned when the PIR-value was 1 in *≥* 900 cell lines, resulting in a fairly balanced dataset (*n_Retained_* = 10635, *n_Spliced_* = 13086).

**Task description** Training sequences were generated by extracting 400 bases before the 5*^′^* and after 3*^′^* intron ends, including 100 bases of the flanking exons to capture potential exonic splice enhancers (ESEs) and silencers (ESSs), resulting in sequences 400 nt long on both sides. This was done as most splice regulatory elements are reported to lie within these limits [89, 90]. Fasta sequences were produced based on genomic coordinates with the Python library pysam (https://github.com/pysam-developers/pysam). Sequences on the negative strand were reverse complemented. The resulting set of intronic sequences were one-hot encoded into four nucleotide channels (A, C, G, T) and forward-passed through the model’s body. Introns shorter than the full 400 base window were padded to 400 bases, with valid positions masked as 1 and padding as 0.

For this task, we used frozen Parnet embeddings extracted from the core pretrained model, computed separately for sequences surrounding the 5*^′^* and 3*^′^* splice sites. The resulting embeddings from both intronic ends were concatenated, mean-pooled, and passed through a shallow two-layer multilayer perceptron (MLP) to predict the probability of intron retention. Model performance was evaluated using the area under the receiver operating characteristic curve (AUROC) and the area under the precision-recall curve (AUPRC), both computed per class in a ten-fold cross-validation setting.

**Comparison with other models** We compared Parnet, using the exact same input sequences, against three baselines: (i) a naive logistic regression classifier trained solely on intron length and GC content, (ii) SpliceAI, and (iii) the general-purpose RNA language model RiNALMo. For RiNALMo and SpliceAI, embeddings from the two intronic ends were extracted in the same manner as for Parnet, concatenated, mean-pooled, and passed through a shallow two-layer multilayer perceptron (MLP) to generate the final prediction. In the case of SpliceAI, we used the model variant of the OpenSpliceAI implementation with a 400 bp receptive field, accepting input sequences of 400 nt. The original nucleotide-level output layer, which produces multi-class probability scores for each position, was replaced with a binary classification head using a sigmoid activation function to predict the retained/non-retained state. In both RiNALMo and Parnet settings, the backbone model weights remained frozen during training, and only the weights of the downstream classifier were updated.

Also for the benchmark methods, model performance was evaluated using AUROC and AUPRC under a ten-fold cross-validation scheme.

### Variant impact scoring

#### Scoring of splicing variants from MutspliceDB

**Datasets** From the MutSpliceDB database ([80] we retrieved 457 curated single-nucleotide variants with experimentally documented effects on pre-mRNA splicing. Each entry comprised a gene symbol, HGVS cDNA notation, allele registry identifier, genome version (hg19 or hg38), and a literature reference.

Genomic coordinates and alleles were derived from the Human Genome Variation Society (HGVS) cDNA notation for each variant (e.g. c.2034+1G>T), leading to one variant being dropped because of its unparseable HGVS notation. Of the remaining 456 variants, 378 were already provided in hg38 coordinates, while the remaining 78 were lifted from hg19 to hg38 using liftover, of which one failed and was discarded. Seventeen variants mapping to non-autosomal chromosomes (chrX, chrY, chrM, or unplaced contigs) were excluded, retaining only variants on autosomes 1–22. Mutations were mapped to GENCODE canonical transcripts, matching by symbol and chromosome, enabling strand assignment. REF alleles were verified against the hg38 reference genome sequence; for variants on minus-strand genes, if the reported REF allele matched the reverse complement of the hg38 base, both alleles were flipped to the forward-strand convention. Five variants for which the reported REF allele could not be reconciled with the reference sequence were discarded. All remaining variants fell within 100 nt of a splice site, and thus each variant was then assigned to its nearest annotated splice site (donor or acceptor). This preprocessing pipeline yielded 433 variants (428 unique HGVS notations) across 338 unique splice sites. Clinical significance was successfully retrieved for 337 of the 428 unique HGVS notation by querying the NCBI E-utilities API (esearch and esummary endpoints), rate-limited to three requests per second in accordance with NCBI usage policy. Clinical significance, star review rating, associated disease name, and linked PubMed identifiers were extracted and cached. Eight variants annotated as having no splicing effect in the source database were excluded from performance evaluation, leaving 425 true-positive variants.

Common genetic variants in the same genomic neighborhoods were extracted from the GnomAD v3.1 ([91]) allele-frequency-only VCF (hg38) as negative controls. Query intervals were defined as *±*50 nt around each MutSpliceDB position, yielding 346 non-redundant intervals. Single-nucleotide substitutions were retrieved, excluding the MutSpliceDB positions themselves and any indels or multi-allelic records. This yielded 2,540 GnomAD SNVs used as candidate negative examples, of which 2,473 were retained as true-negative controls after restricting to those co-localized with the 425 retained positive variants.

We additionally introduced an *in silico mutagenesis* (ISM) analysis over 101-nt windows (splice site position *±* 50 nt) centered at each of the 338 unique splice sites. At every position within each interval, all three possible single-nucleotide substitutions were enumerated relative to the hg38 reference allele, generating a complete (ISM) map that enabled ranking of individual MutSpliceDB mutations within their local background.

**Task Description** Parnet was applied to score each variant by comparing predictions for the reference and alternate allele sequences, each provided as a 601-nt window centered on the variant position. Impact was measured per RBP from the binding profiles by either measuring the Jensen-Shannon Divergence (JSD) for each RBP binding profile (computed over a central 101 nt sub-interval to measure widespread alterations of binding), or calculating Δ*P*, the difference in cumulative probability over a local *±*5 nt interval centered on the mutated position (measuring predicted increase or decrease in local binding). Aggregation (average or maximum) was done either from all 223 RBP tracks, or from the subset of 81 splicing-related RBPs combined with the core set of 27 spliceosomal RBPs. Besides binding profiles, embeddings were used to score mutations by computing the cosine distance between reference and alternate sequence representations extracted from the Parnet encoder, computed at each individual position, and averaged for the *±* 5nt centered interval. Performance was evaluated in the context of a binary classification task, computing Receiver operating characteristic (ROC) and precision-recall (PRC) curves, and reporting area under curves for each scorer.

**Comparison with other models** For comparison we considered the OpenSpliceAI [92] implementation of the SpliceAI model [12], using the dedicated scoring tools. The model takes a 10, 000-nt genomic sequence centered on the variant as input and predicts, at each output position, the probability of splice donor gain (DG), donor loss (DL), acceptor gain (AG), and acceptor loss (AL). For each variant, delta scores were computed as the difference in predicted probabilities between the alternate and reference sequences, and DS_max = max(Δ*DG*, Δ*DL*, Δ*AG*, Δ*AL*) was evaluated at the variant position (distance parameter set to 1, masking disabled). We used the ensemble of five independently trained models (rs10–rs14) provided by authors, and worked with the averaged predictions as ouptut by the OpenSpliceAI scoring script. We note that for 121 of 2,898 variants (11 MutSpliceDB positives and 110 matched gnomAD controls, localized to the CHD2, MGA, PRKDC, and RECQL4 loci), OpenSpliceAI returned no delta score because these genes were absent from the transcript annotation supplied to the tool. This is a limitation of OpenSpliceAI-family predictors, requiring an annotation to anchor their donor/acceptor probability tracks, while other models work annotation-free. OpenSpliceAI’s ROC/PRC in this comparison is therefore restricted to the remaining n=2,777 variants.

We also included RiNALMo as a reference pre-trained RNA language model to compare embedding-based scoring. As with Parnet embeddings, each variant was scored by computing the cosine distance between the encoder representations of the reference and alternate nucleotides within input 601-nt sequences, and averaging for the central 11 nucleotides. We also computed the cosine distance from the embedding representation of the *CLS* token produced by RiNALMo as a proxy of the overall sequence.

Finally, Nucleotide-level conservation scores were obtained from the UCSC 100-way vertebrate phyloP track [93]. Positive scores indicate evolutionary constraint relative to the expected neutral rate; negative scores indicate accelerated evolution. The PhyloP score at the variant position was used directly as a discriminator of MutSpliceDB true positives from GnomAD background controls.

### Scoring SpliceBench Splicing Reporter Assays mutations

**Datasets** Six functional variant effect datasets compiled by [94] were used for evaluation.

One dataset was generated by saturation genome editing (SGE) using homology-directed repair (HDR) in HAP1 cells: BRCA1, covering 11 exons (1,386 variants). The remaining five datasets were generated by minigene splicing reporter assays: FAS exon 6 (189 variants), MLH1 spanning 18 exons (296 variants), POU1F1 exon 2 (941 variants), RON exon 11 (598 variants), and WT1 exon 9 (502 variants), for a total of 3,912 variants across six genes. Strand orientation was assigned from the GENCODE canonical transcript for each gene, and REF alleles were verified against the hg38 reference sequence at reported positions. For downstream analysis, we considered the reported functional scores and the “Splicing-Disrupting Variant” (SDV) label pre-assigned by the original authors. To explore the functional landscape of the different regions, we introduced an *in silico mutagenesis* analysis, as done above, for each of the datasets by identifying exon boundaries from the GENCODE canonical transcript of each assayed exon, extending their intervals to 50 nt into each flanking intronic sequence.

**Task Description** As for the MutSpliceDB analysis, we applied different scoring methods using Parnet to predict the functional impact of mutations. Using the Binary SVD labels, we computed Receiver operating characteristic (ROC) and precision-recall (PRC) curves, reporting area under curves for each scorer, either across datasets or per dataset. Individual mutations were further characterized through the binding impacts as measured by the difference in cumulated binding probabilities, with respect to local GnomAD mutations as well as ISM profiles.

**Comparison with other models** We considered the precomputed SpliceAI scores. We also included RiNALMo as a reference pre-trained RNA language model to compare embeddings based scoring, applying the same approach as described for the MutSpliceDB analysis.

Splice-altering potential was additionally scored with OpenSpliceAI (variant subcommand), using a 5-model ensemble of OpenSpliceAI-MANE checkpoints (10,000 nt flanking context; random-seed replicates rs10–rs14), whose predicted reference/alternate splice-site probability tracks were averaged across the 5 models before computing delta scores. Scoring used the GRCh38/hg38 reference genome and a GENCODE-derived gene annotation, with –distance 1 (restricting the donor/acceptor gain-loss search window to the variant position itself and direct neighbouring positions) and –mask 0 (no masking of annotated-gain/unannotated-loss scores). For each variant we report DS_max, the maximum of the four raw delta scores (donor gain/loss, acceptor gain/loss) across all overlapping gene annotations. The same configuration was applied identically to MutSpliceDB and SpliceBench variants. Finally, SNVs were annotated with the PhyloP 100w score, using hg38 phyloP100way (UCSC hg38.phyloP100way.bw) at the single mutated base position, via pyBigWig. This measures evolutionary conservation across 100 vertebrates, and grouped so as to separate highly constrained genomic positions (positive PhyloP scores) from neutrally or fast evolving positions (negative PhyloP scores).

### Additional analysis

#### RNA splicing maps

For RNA maps analysis exon sets derived from siRNA knockdown RNA-Seq were integrated from several sources (Supplementary Table 1). For eCLIP, Skipper “finemapped” 75nt peaks were retrieved from Figshare [95]. To generate comparable Parnet peaks, Parnet ‘target’ predictions along MANE v1.4 full transcripts (restricted to the top 50% expressed transcripts) were lifted to GRCh38 genomic coordinates. Parnet per-base predicted-binding scores were scaled *×* 1000 and rounded to integers; therefore positions with a raw predicted score below 5 *×* 10*^−^*^4^ (rounding to zero) were discarded. Peaks were called using Clippy settings -n 15 -w 0.8 -x 3 -mx 5 -mb 5 (www.github.com/ulelab/clippy). Then, to match the fixed-width Skipper peaks, each Clippy peak was reduced to its midpoint and extended by 37 nt on each side to give a 75-nt window. RNA maps were generated with www.github.com/ulelab/rna_maps (VastDB mode on branch feat-bootstrapping), which computes the positional enrichment of protein binding around regulated exons relative to a control exon set. The per-exon *×* per-position coverage matrix was binarised to 0/1 (each cell records whether the exon had at least one overlapping binding feature at that base), so that the per-position signal for a category is the fraction of exons bound at that position. Enrichment was quantified with a bootstrap contrast (--enrichment bootstrap_contrast): for each regulated category *c* (enhanced or silenced) versus control, exons were resampled with replacement (*B* = 1000 iterations, --n_boot 1000, seed 42), and at each position we report

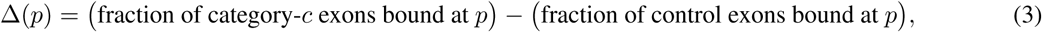

together with its 2.5–97.5th percentile bootstrap confidence interval as a shaded ribbon underneath.

In cases where multiple RNA-seq datasets were available upon knockdown of the same RBP, we choose the dataset that generated the RNA map with the highest signal when using either eCLIP or Parnet predictions. More in detail, to present a single, most-informative map per protein we ran all possible combinations and then extracted the peak signal as the position of largest |Δ| within the enhanced and silenced categories, and scored it by the bootstrap-CI bound on |Δ| closest to zero (i.e. the largest effect defensible at the 95% CI level; set to zero where the CI straddled zero). Per protein *×* exon source *×* cell line, the combination score was the maximum of this CI-bounded peak across eCLIP and Parnet and across the enhanced/silenced categories. For each protein we selected the exon source and cell line with the largest combination score; where no combination reached significance on either source, we fell back to the largest point-estimate peak. To filter for RNA maps with meaningful signal above control, we show all maps where peak |Δ| reached *≥* 0.12 in at least one source/category/region.

#### Parnet validation with *in vitro* binding motifs

For a given RBP and celltype eCLIP experiment, we computed feature importance through Integrated Gradients from up to 1,000 highest ranking NarrowPeaks considering their *signalValue* (enrichment of eCLIP signal over SMI signal), considering NarrowPeaks from both replicates. In each sequence, the top scoring 5-mer was identified. By aggregating (summing 5-mer importance scores) across sequences, we generated a table of 5-mer with so-called relevance scores, for each of the 3 tracks.

We correlated these scores with 5-mer scores derived from *in vitro* experiments, namely RNAcompete [53, 54] and RNA-Bind-n-Seq [96], which were processed as described in [32]. To evaluate the denoising impact of the penalized mixing, we compared the Spearman correlations of 5-mer importance scores derived from the target (denoised) track against the correlations of 5-mers from the total track. To assess whether Parnet’s target track more faithfully captures genuine protein-specific binding signals, we compared the agreement between *in vitro* and textitin vivo 5-mer enrichment for the target and control tracks across RBPs. Specifically, we quantified the difference in correlation between in vitro and in vivo 5-mer scores for the target track relative to the control track, where higher values indicate that the target track more closely recapitulates experimentally determined in vitro binding preferences.

This difference in correlations is then calculated systematically from Parnet models trained over a range of increasing penalty factors. The Δ correlations are reported for each RBP, averaging over cell-types when an RBP is measured in both HepG2 and K562. Furthermore, the average value is calculated for each RBP over the range of penalty factors, to identify those that overall benefit from the addition of the *λ* penalty to the model.

#### Sequence Importance Scores with Integrated gradients (IGs)

To identify nucleotide positions that drive Parnet predictions, we computed per-position feature importance scores using Integrated Gradients [71], as implemented in the Captum library [97]. IG attributes a model’s output to each input feature by approximating the integral of gradients along a straight interpolation path from a reference baseline to the actual input. Here, inputs were one-hot encoded sequences of 601 nucleotides (shape *Batch ×* 4 *× Length*), and the baseline was an all-zeros tensor of the same shape, representing a uniform, uninformative sequence. The integral was approximated using 20 interpolation steps. For each sequence and each RBP–cell-type (RBP_CT) task, attributions were computed with respect to the central 101 output positions (*±*50 nucleotides around the sequence midpoint), so that the resulting importance scores reflect each nucleotide’s contribution to predictions in the peak center rather than flanking regions. Per-position importance was derived by multiplying the mean attribution across the 101 target positions by the one-hot encoded input and summing over the four nucleotide channels, yielding a scalar importance value per position. Attributions were computed independently for the total, target and, control output tracks of Parnet. Convergence of the IG approximation was verified by inspecting the convergence delta, which remained below 5 *×* 10^−3^ across all sequences.

We further used the IG-based approach described above to estimate per-nucleotide feature importance for the downstream tasks. The interpolation steps and baseline settings remained the same, i.e. 20 steps and all-zero one-hot encoded uninformative tensor as a baseline. For each input sequence passed through the model, the attributions were computed with respect to the final prediction; in case of the intron retention task, the logit produced by the final joint classifier, in the case of splicing site classification, the class logit for the sequence’s true label, obtained by averaging the position-wise outputs of the classification head over the sequence. Here we report attributions for the donor and acceptor classes. For each class, attributions were computed on the 500 sequences to which the model assigned the highest probability of that class (restricted to sequences whose true label matched), using 50 interpolation steps and the same all-zero baseline as above.

#### Attribution motifs

To identify the sequence features underlying the model’s attribution scores, we applied tfmodisco-lite, a motif discovery algorithm that identifies recurring patterns in deep-learning attribution maps, hereafter referred to as ‘attribution motifs’. For all downstream tasks, we followed the standard procedure described in the package. From all inferred attribution motifs, we retained only those supported by at least 20 seqlets (i.e., sequences assigned to a motif) at an FDR of 0.05. These motifs were matched with TomTom against a comprehensive collection of human RBP motifs [54]. For visualization, we report the top three RBP motif matches with a TomTom q-value below 0.05.

#### Embedding analysis and visualization

To visualize learned representations, embeddings were extracted from the penultimate layer of the Parnet model and projected into two dimensions using UMAP dimensionality reduction. The embedding representation and visualization strategy were adapted to each downstream task. Unless otherwise specified, UMAP was applied to full-dimensional mean-pooled embeddings using the default hyperparameters (*n*_neighbors_ = 15, min_d_ist = 0.1).

For the RNA families and genomic regions classification tasks, separate UMAP projections were generated for each dataset, with all sequences pooled into a single shared embedding space per task. The resulting projections were first colored by class label to evaluate separation between RNA biotypes or genomic regions. The same UMAP coordinates were subsequently colored by GC content and transcript length to assess the relationship between sequence-level covariates and embedding geometry.

For the lncRNA localization task, all transcripts were projected into a shared two-dimensional embedding space and initially colored by dissociation class (fast vs. slow) to assess separation between chromatin-retained and nucleoplasmic lncRNAs. The same projection was then colored by half-life quartiles, binding affinities of individual RBPs (SNRNP70, XRN2, SRSF1, SRSF7, SRSF9), as computed in Ntini et al. [56], transcript length, and transcribed enhancer histone marks (H3K4me1, H3K4me3) to identify biological features associated with embedding structure.

To examine how translational output relates to the Parnet embedding space, we examined two-dimensional embeddings of PC3 transcripts. Transcripts were colored by their log(TE) quartile (bottom 25% vs. top 25%) to evaluate whether highly and weakly translated transcripts occupy distinct regions of the embedding space. To further relate embedding structure to predicted regulatory interactions, RBP binding-site densities were computed from nucleotide-resolution Parnet binding probability profiles. For each transcript and RBP, binding peaks were identified as positions exceeding the empirical background probability distribution (empirical *p*-value threshold), and significant positions within 5 nucleotides were merged into a single peak. Peak counts were normalized by transcript length to obtain RBP-specific binding-site density, which was used as a proxy for overall binding occupancy. The UMAP projection was subsequently colored by binding-site density for translation-associated RBPs (e.g. LARP4, EIF3D, EIF4G2).

For intron retention prediction, embeddings corresponding to the 5*^′^* end, middle, and 3*^′^* end of each transcript were average-pooled separately and concatenated before UMAP projection. The resulting embedding space was colored by percentage of intron retention (PIR), highlighting transcripts within the highest and lowest 25% of PIR values.

For splice-site prediction, no pooling was performed, and nucleotide-level embeddings were projected directly using UMAP. In this case, UMAP was run with modified parameters (*n*_neighbors_ = 50, cosine distance metric, min_dist = 0.15). The resulting embeddings were colored by splice-site classification label (donor, acceptor, or neither) to evaluate the separation of nucleotide-level representations by functional class.

#### Mutation impact interpretation

For each mutation, we computed the difference of binding probability of each of the 223 RBP-cell-type tracks between the mutated sequence and the reference sequence. This provides with a signed score that quantifies the predicted increased or decreased binding induced by the mutation. These measurements were used to cluster mutations and RBPs, as well as to identify the most impacted RBPs for individual mutations of interest. When applicable, mutation impact was further illustrated by computed attribution scores through integrated gradients, highlighting RBP motifs and how the mutation affects them. Finally, in the context of classification task and local background mutations, mutations of interest were characterized relative to background mutations in their vicinity, either from the ISM results or from datasets specific negative control mutations.

## Declarations

### Ethics approval and consent to participate

Not applicable.

### Consent for publication

Not applicable.

### Availability of data and materials

Code for Parnet model training and applications is available at https://github.com/marsico-lab/parnet. Code for the analyses presented in this paper is available at https://github.com/marsico-lab/parnet--paper Code in both repositories is available under the license Apache License 2.0.

All data processed in this study was obtained exclusively from public sources. CLIP-seq data was obtained from ENCODE [50] (eCLIP). *In vitro* data on protein-RNA interaction was obtained from Dominquez et al. [96] (RNA-Bind-n-Seq) and Ray et al. [53] (RNAcompete). Splicing-related mutations were obtained from MutSpliceDB [80] and SpliceBench [82], while further mutations were obtained from gnomAD [91]

### Competing interests

The authors declare no competing interests.

## Acknowledgments

We acknowledge Ole Winther and Jun Wang, University of Copenhagen for fruitful discussions. We thank Marcel Schulz, Goethe University Frankfurt, and Kathi Zarnack, University of Würzburg, for their valuable discussions, insightful suggestions, and critical reading of the manuscript. We acknowledge the use of the HPC cluster at Helmholtz Munich for the computational resources used in this study. We acknowledge the use of ChatGPT and Grammarly for text editing.

## Funding

This work was supported by the Helmholtz Association under the joint research school “Munich School for Data Science (MUDS)” to A.M., A.T. and S.L. It was further supported by the BMBF Cluster4Future program (Cluster for Nucleic Acid Therapeutics Munich, CNATM, ID: 03ZU1201AA) to A.M. amd A.T., SFB TRR267 (Project-ID 403584255) to A.B. and A.M., the Helmholtz Association’s Initiative and Networking Fund through “RBPAI4Virus” (grant # ZT-I-PF-5-176) to L.M. and the Deutsche Forschungsgemeinschaft (DFG) – Project-ID 533767322 – EXC 3113/1, Cluster for Nucleic Acid Sciences and Technologies – NUCLEATE to A.M. G.D. was supported by the DFG (TRR 267/A09, 403584255; Basic Module, 562550297) and Germany’s Excellence Strategy (CPI, EXC 2026, 390649896; SCALE, EXC 3094, 533751785). In addition, we acknowledge the European Research Council under the European Union’s Horizon 2020 research and innovation programme (835300-RNPdynamics to J.U.), the Wellcome Trust (227465/Z/23/Z to J.U.), by The Francis Crick Institute, which receives its core funding from Cancer Research UK (CC0102), the UK Medical Research Council (CC0102), the Wellcome Trust (CC0102) and the UK Dementia Research Institute (award number UK DRI-RE21605), principally funded by the UK Medical Research Council.

## Authors’ contributions

M.H. and A.M. conceived the project. M.H. collected and processed the datasets, conceptualized and implemented the PARNET model; L.M and A.T. led and performed most of the analysis and Parnet fine-tuning downstream tasks; A.B., C.K. and K.K. contributed functional analysis on intron retention, functional exons and sequence motifs. J.G. and G.D. contributed to the conceptualization of the intron retention and mutation analysis tasks. J.U. supervised and offered insights into the motif, spliced exon, and intorn retention analysis. M.H. and A.M supervised and guided the project, with inputs from J.U.; AM provided resource, compute infrastructure and wrote the manuscript with help from M.H., L.M., A.T., A.B., C.K. and J.U; All authors reviewed and approved the final manuscript.

## Notes

https://github.com/marsico-lab/parnet--paper

### Competing Interest Statement

The authors have declared no competing interest.

