## Supplementary material for "PARNET: A CLIP-SEQ-BASED FOUNDATION MODEL FOR RNA SEQUENCE REPRESENTATION LEARNING"

### Supplementary Figures

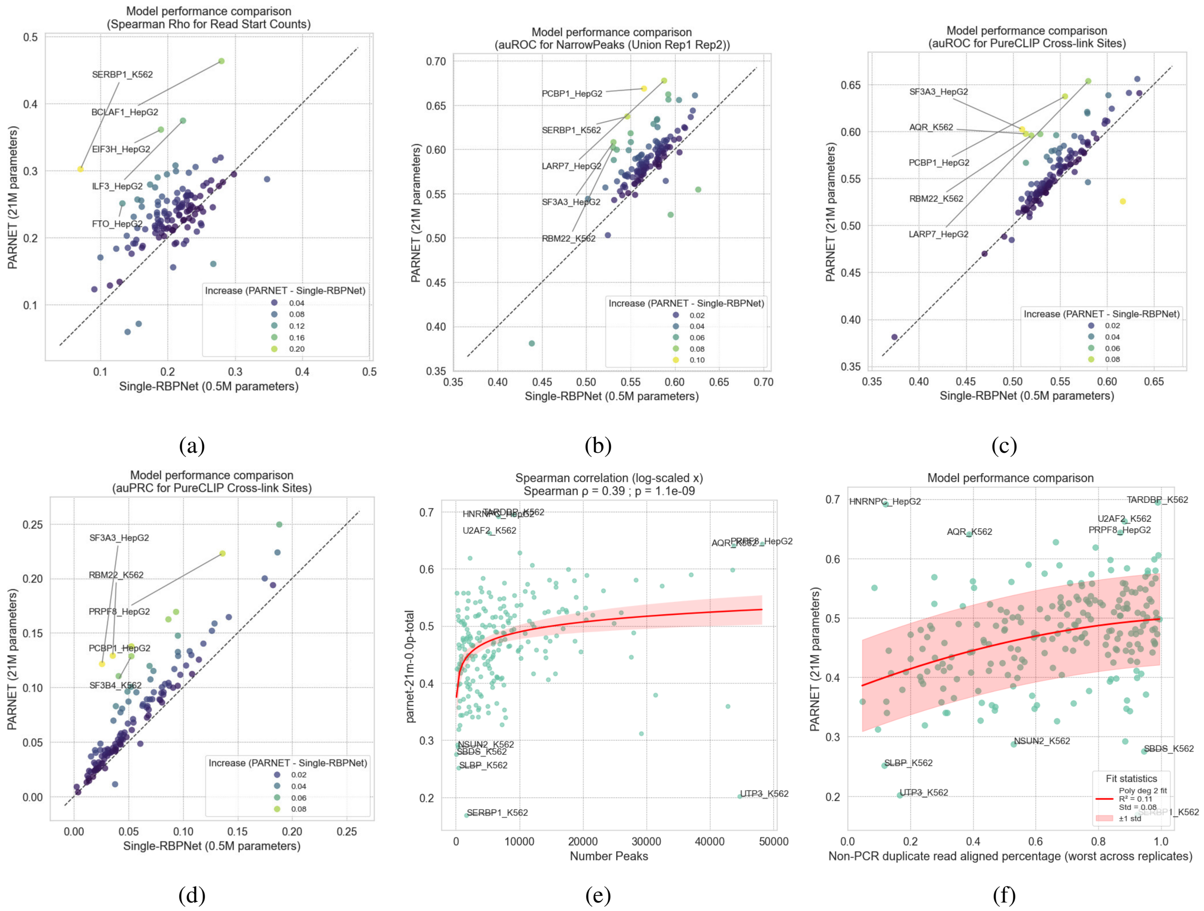


Supplementary Figure 1: Extended Data Figure 1 (related to main Figure 2). Additional evaluations of PARNET performance on eCLIP datasets. **a-d** Performance of the PARNET model, measured by Pearson correlation, plotted against the cross-replicate Pearson correlation of ground-truth read-start counts derived from the original CLIP datasets. **e-f** Relationship between PARNET performance of individual RBP predictions and proxies of datasets quality, as measured from the number of called NarrowPeak (e) or from the percentage of non-PCR-duplicate aligned reads (f).

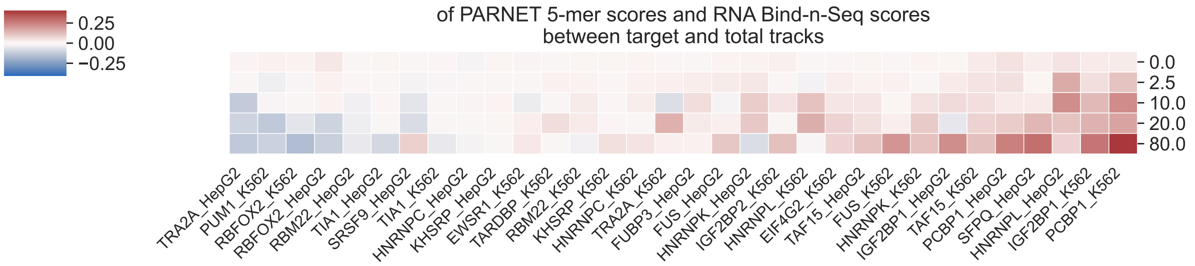


(a)

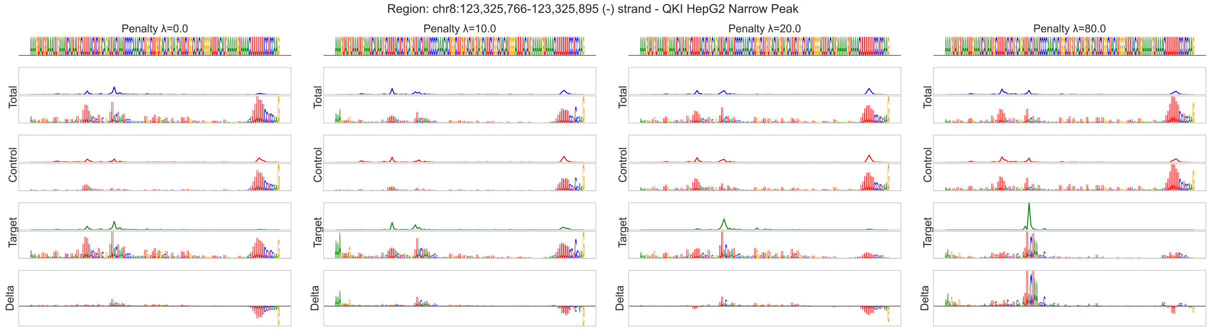


(b)

Supplementary Figure 2: Extended Data Figure 2 (related to main Figure 2). Additional motif evaluation. **a** (Related to main Figure 2e). Difference ($\Delta$) in correlation between *in vitro* 5-mer scores (from Bind-n-Seq assays) and PARNET feature importance across increasing penalty. For each RBP, $\Delta$ is computed as the difference between correlations obtained from the denoised target and total prediction tracks. Higher $\Delta$ indicates improved agreement between *in vivo* and *in vitro* motifs and more faithful recovery of genuine target motifs. **b** (Related to Figure 2d). Additional example of PARNET predicted track and motif denoising at increasing $\lambda$ penalty, here on a binding region of QKI in cell-type HepG2.

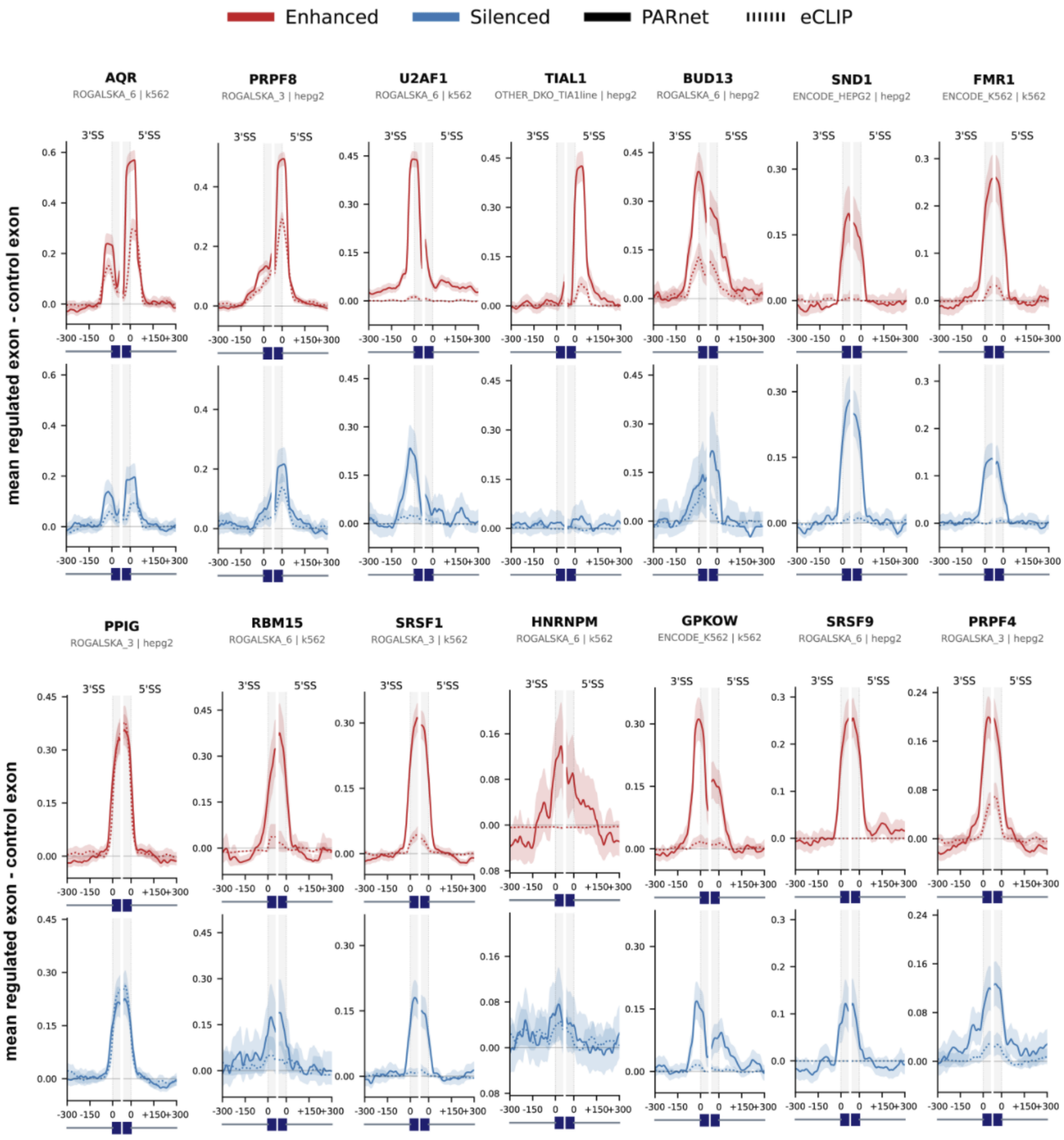


(a)

Supplementary Figure 3: Extended Data Figure 3 - related to main Figure 2g. Exon–intron meta-profiles of RBP binding estimated from eCLIP signal (dotted lines) and PARNET predictions (solid lines) across four classes of splicing deregulation following knockdown of an expanded set of RBPs

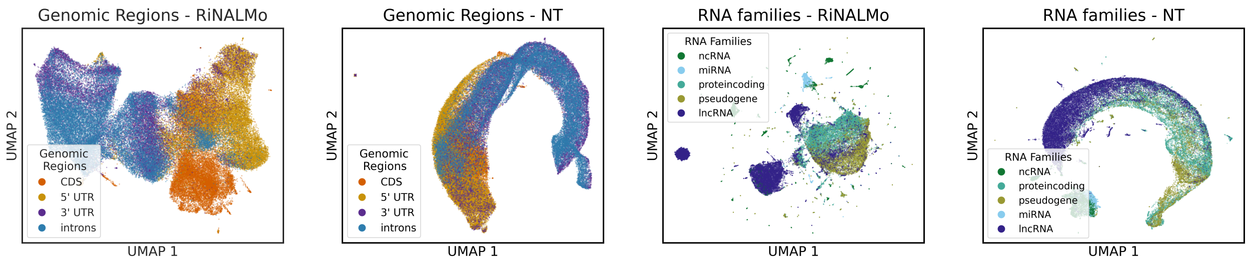


(b) (c) (d) (e)

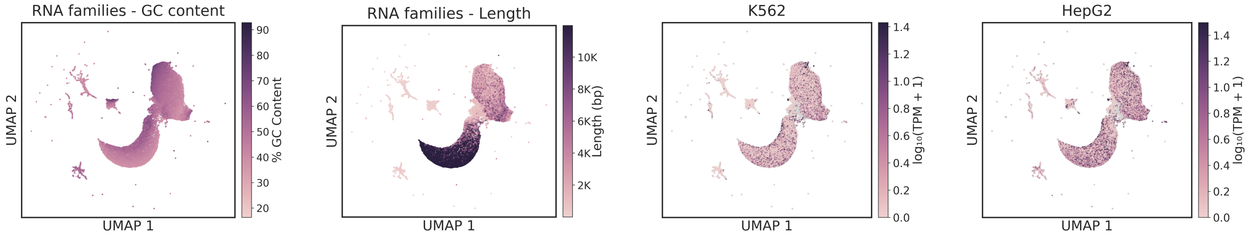


(f) (g) (h) (i)

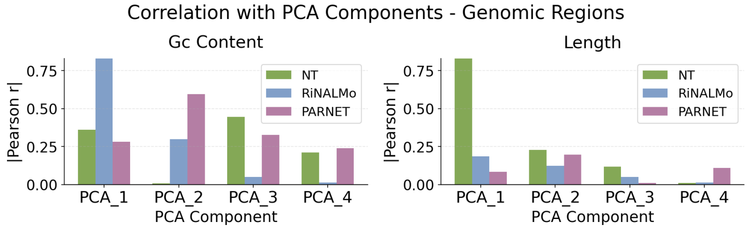


(j)

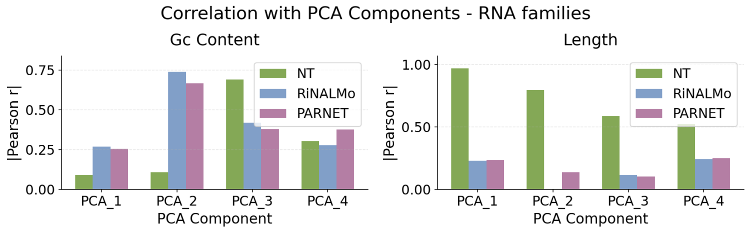


(k)

Table 1: Genomic Regions: Per-component explained variance ratio (%)

| Component | NT | RiNALMo | Parnet |
|---|---|---|---|
| PCA 1 | 88.19 | 24.52 | 37.69 |
| PCA 2 | 5.01 | 13.74 | 21.56 |
| PCA 3 | 1.29 | 6.95 | 6.06 |
| PCA 4 | 0.81 | 5.53 | 5.56 |

Table 2: RNA Families: Per-component explained variance ratio (%)

| Component | NT | RiNALMo | Parnet |
|---|---|---|---|
| PCA 1 | 81.48 | 22.53 | 78.93 |
| PCA 2 | 7.04 | 16.48 | 6.22 |
| PCA 3 | 2.38 | 10.82 | 3.11 |
| PCA 4 | 1.00 | 9.23 | 2.12 |

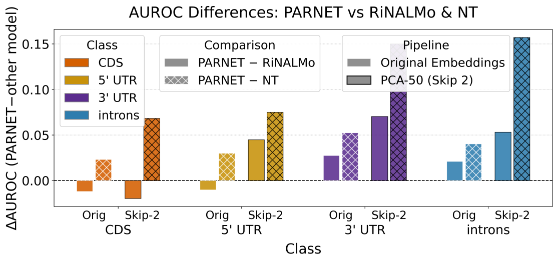


(l)

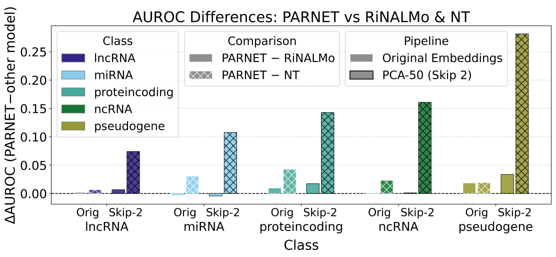


(m)

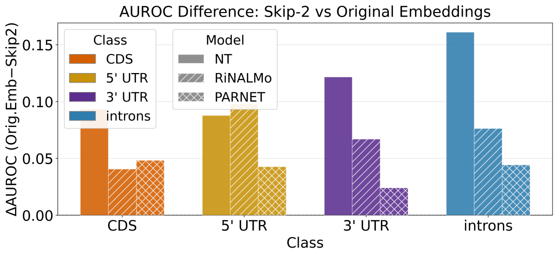


(n)

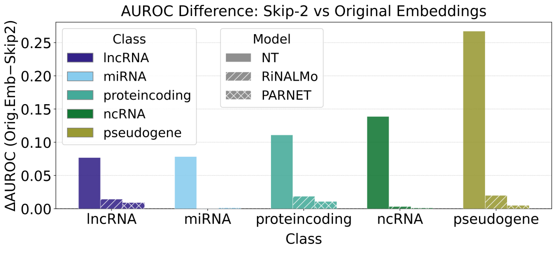


(o)

Supplementary Figure 4: Supplementary Figure 4

**Additional Data Figure 4 - Related to Figure 2b-f** **(a)** UMAP of RiNALMo embeddings for GENCODE transcripts, colored by genomic element. **(b)**UMAP of Nucleotide Transformer embeddings for GENCODE transcripts, colored by genomic element. **(c)** UMAP of RiNALMo embeddings for GENCODE transcripts, colored by RNA family. **(d)** UMAP of Nucleotide Transformer embeddings for GENCODE transcripts, colored by RNA family. **(e)** UMAP of PARNET embeddings of RNA families, colored by GC content. **(f)** UMAP of PARNET embeddings of RNA families, colored by transcript length. **(g)**UMAP of PARNET embeddings of RNA families, colored by normalized transcript expression in K562 ($\log_{10}(\mathrm{TPM})$). **(h)**UMAP of PARNET embeddings of RNA families, colored by normalized transcript expression in HepG2 ($\log_{10}(\mathrm{TPM})$). **(i)**Correlation of the first four principal components (PC1 - PC4) of the PARNET, RiNALMo, and Nucleotide Transformer genomic region embeddings with GC content (left) and sequence length (right). The accompanying table (Table 1)reports the proportion of variance explained (%) by each principal component. **(j)** Correlation of the first four principal components (PC1 - PC4) of the PARNET, RiNALMo, and Nucleotide Transformer RNA families embeddings with GC content (left) and sequence length (right). The accompanying table (Table 2)reports the proportion of variance explained (%) by each principal component. **(k)** Difference in performance ($\Delta\mathrm{AUROC}$) between PARNET and either RiNALMo or the Nucleotide Transformer on the genomic region classification task, reported for each class using both the original embeddings (*Orig*) and embeddings with the first two principal components removed (*Skip-2*) **(l)** Difference in performance ($\Delta\mathrm{AUROC}$) between PARNET and either RiNALMo or the Nucleotide Transformer on the RNA family classification task, reported for each class using both the original embeddings (*Orig*) and embeddings with the first two principal components removed (*Skip-2*) **(m)** For each model (PARNET, RiNALMo, and Nucleotide Transformer), the decrease in performance on the genomic region classification task is reported as $\Delta\mathrm{AUROC}$ between the full embeddings and embeddings with the first two principal components removed. **(n)** For each model (PARNET, RiNALMo, and Nucleotide Transformer), the decrease in performance on the RNA family classification task is reported as $\Delta\mathrm{AUROC}$ between the full embeddings and embeddings with the first two principal components removed.

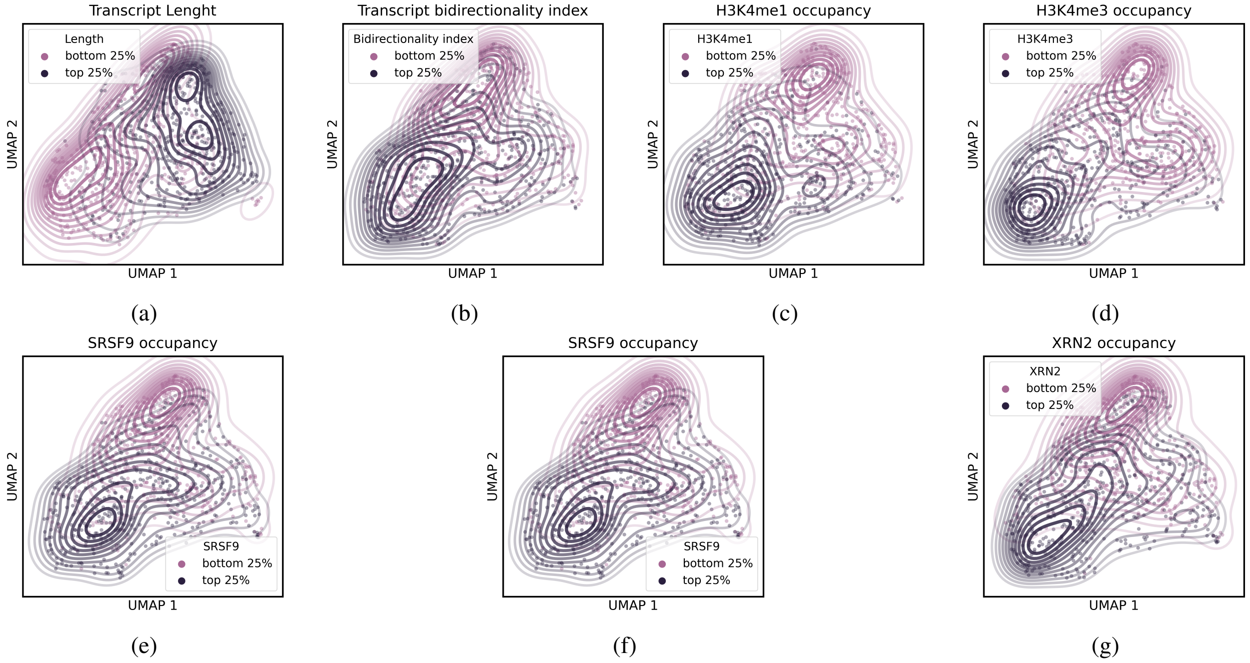


Supplementary Figure 5: Additional Data Figure 5 - Related to main Figure 3g-l. Parnet embedding space colored by transcript length (a), transcript bidirectional index (b), H3K4me1 occupancy (c), H3K4me1 occupancy (d), SRSF9 ENCODE peaks occupancy (e), SRSF9 ENCODE peaks occupancy (f) and XRN2 NCODE peaks occupancy (g), with contour lines marking the upper and lower 25% quantiles of the occupancy distribution.

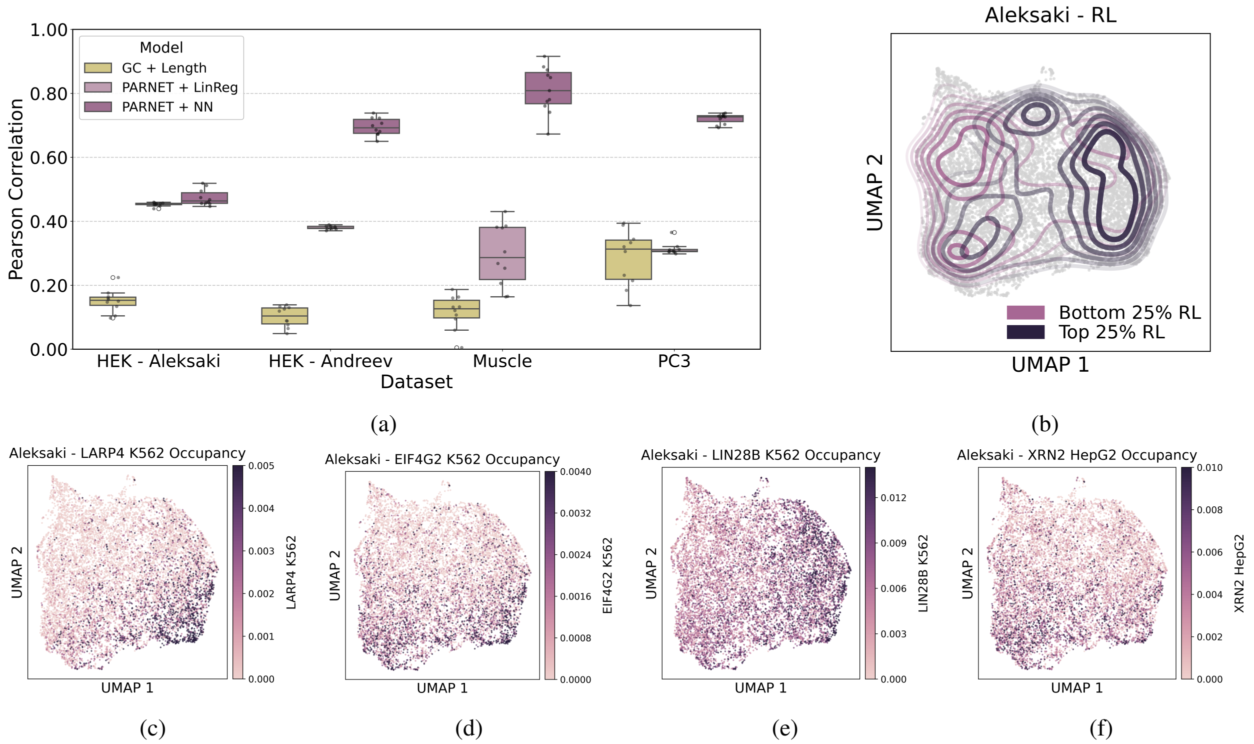


Supplementary Figure 6: (a) Predictive performance (Pearson correlation) of translational efficiency (TE) models using either GC content and transcript length as baseline covariates, or Parnet-derived embeddings with two downstream approaches: a classifier and a neural network model. Error bars represent variability across folds in a 10-fold cross-validation scheme evaluated on four independent datasets. (b) Two-dimensional UMAP projection of the Parnet embedding space colored by TE values in the Aleksaki HEK293 dataset (for comparison to the PCs dataset in Figure 4). Contour lines indicate the upper and lower 25% quantiles of the TE distribution. (c–f) Two-dimensional UMAP projections of the Parnet embedding space colored by RNA-binding protein occupancy for LARP4 (c), EIF4G2 (d), LIN28B (e), and XRN2 (f).

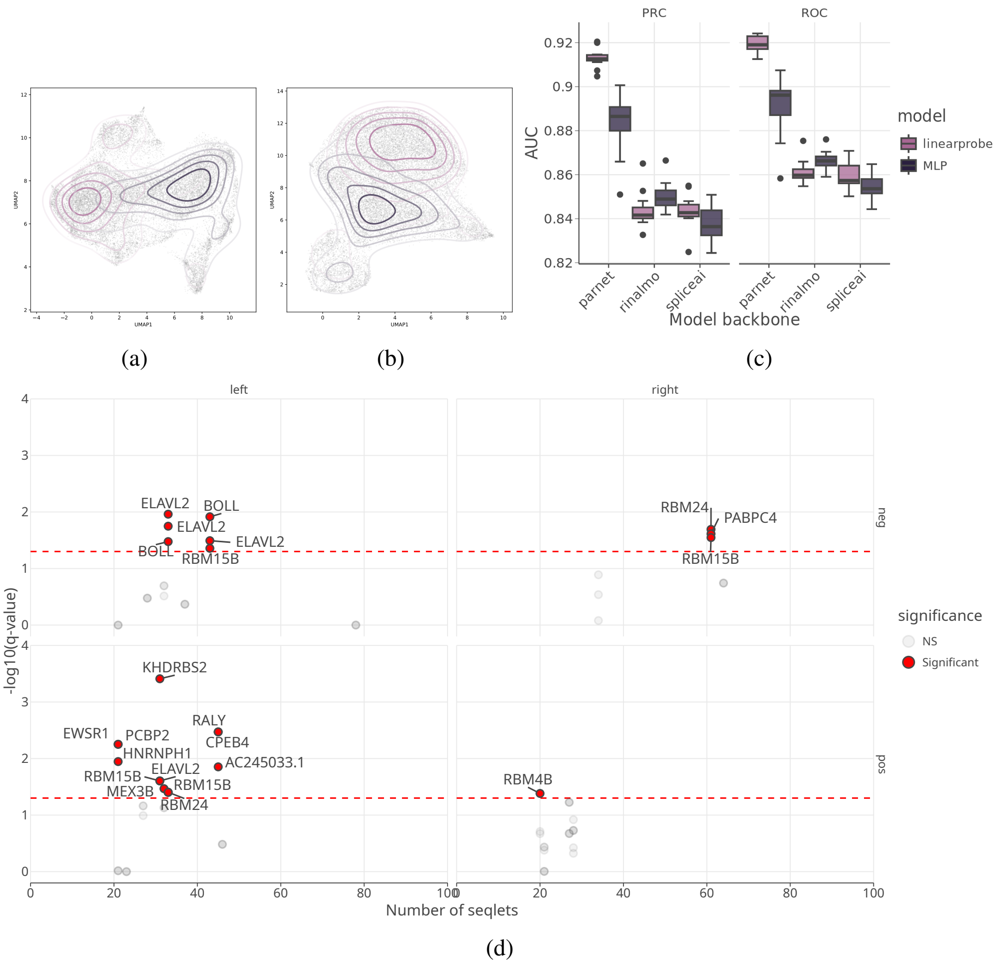


Supplementary Figure 7: Prediction of intron retention with PARNET. **a-b** Two-dimensional UMAP projection of the RinAlmo (a) and SpliceAI (b) embedding space, where each points represents an intron in the dataset. Contour lines indicate the upper (dark purple) and lower (pink) 25% quantiles of the TE distributionRetained (dark purple). **C** Boxplots showing area under PRC (left) and ROC (right) curves for 10 CV folds, comparing a linrar classifier versus an MLP model on frozen embeddings for Parnet, RinAlmo and spliceAI. **D** Results of TomTom analysis of motifs extracted from IG attributions by tf-modisco. Each dot corresponds to the IG-derived motif. Names of the top matching RBP motifs from the euPRI database are reported directly on the plot, with significance level plotted on Y and the number of motif occurrences on X. Dotted red lines correspond to significance levels.

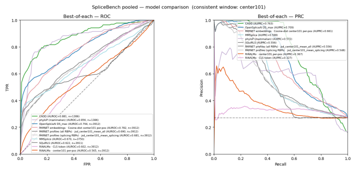

(a)

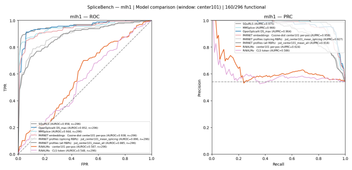

(b)

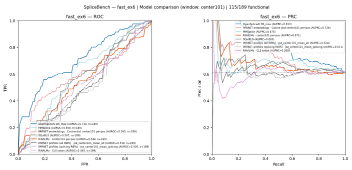

(c)

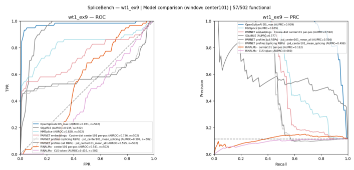

(d)

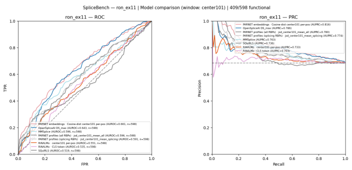

(e)

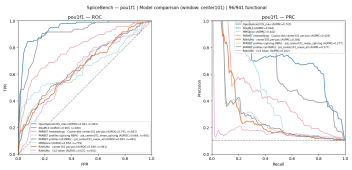

(f)

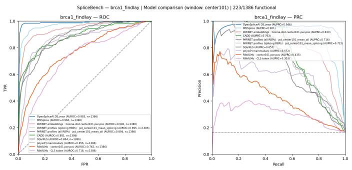

(g)

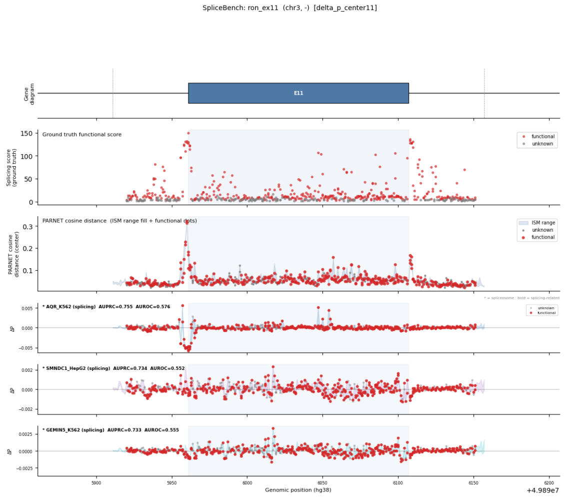


(h)

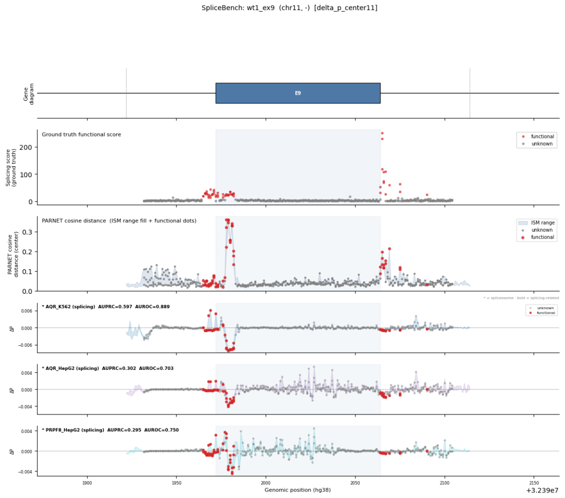


(i)

Supplementary Figure 8: Additional results on mutations analyses. **(a)** ROC and PRC on SpliceBench mutations, aggregated over the different datasets. **(b-g)** ROC and PRC on SpliceBench mutations, for each of the different datasets. **(h,i)** Examples of two profiles, showing the ground-truth scores and their classification, the PARNET embeddings-derived scores, and the top 3 RBPs with highest area under the PRC.

### Supplementary Tables
